# Dominant species determine ecosystem stability across scales in Inner Mongolian grassland

**DOI:** 10.1101/2021.10.31.466650

**Authors:** Yonghui Wang, Shaopeng Wang, Liqing Zhao, Cunzhu Liang, Bailing Miao, Qing Zhang, Xiaxia Niu, Wenhong Ma, Bernhard Schmid

**Affiliations:** Ministry of Education Key Laboratory of Ecology and Resource Use of the Mongolian Plateau & Inner Mongolia Key Laboratory of Grassland Ecology, School of Ecology and Environment, Inner Mongolia University, Hohhot, 010021, China; Institute of Ecology, College of Urban and Environmental Sciences, and Key Laboratory for Earth Surface Processes of the Ministry of Education, Peking University, Beijing, China; Department of Geography, Remote Sensing Laboratories, University of Zürich, Winterthurerstrasse 190, 8057 Zürich, Switzerland

**Keywords:** Biodiversity, Productivity, Scale dependence, Species synchrony, Precipitation, Climate change

## Abstract

There is an urgent need to extend knowledge on ecosystem temporal stability to larger spatial scales because presently available local-scale studies generally do not provide effective guide for management and conservation decisions at the level of an entire region with diverse plant communities. We investigated temporal stability of plant biomass production across spatial scales and hierarchical levels of community organization and analyzed impacts of dominant species, species diversity and climatic factors using a multi-site survey of Inner Mongolian grassland. We found that temporal stability at a large spatial scale, i.e. a large area aggregating multiple local communities, was related to temporal stability of and asynchrony among spatially separated local communities and large-scale population dynamics of dominant species, yet not to species richness. Additionally, a lower mean and higher variation of yearly precipitation destabilized communities at local and large scales by destabilizing dominant species population dynamics. We argue that, for semi-arid temperate grassland, dynamics and precipitation responses of dominant species and asynchrony among local communities stabilize ecosystems at large spatial scales. Our results indicate that reduced amounts and increased variation of precipitation may present key threats to the sustainable provision of biological products and services to human well-being in this region.

## Introduction

The ability of ecosystems to stably provide biological products and services such as biomass production for human well-being (Isbell et al., 2015; Tilman et al., 2014, 2006) is being threatened by species loss (Cardinale et al., 2012; Harrison et al., 2015; Isbell et al., 2017, 2015; Newbold et al., 2015; Tilman et al., 2014) and pronounced climatic changes (Hautier et al., 2015, 2014; Ma et al., 2017; Xu et al., 2015). Policymakers seek guidance to make management and conservation decisions at high levels of ecological organization, e.g. an entire region with diverse plant communities (Cardinale et al., 2012; Isbell et al., 2017; Manning et al., 2019; Wang et al., 2019), here referred to a large-scale community (Figure 1a). However, previous theoretical, experimental and observational studies on ecosystem temporal stability have mostly been conducted at local scales with constant environmental conditions (Hautier et al., 2015, 2014; Hector et al., 2010; Isbell et al., 2015; Ma et al., 2017; Tilman et al., 2006; Wang et al., 2020). Patterns of ecosystem temporal stability discovered in local communities may not directly scale up to a system of spatially separate communities (Lamy et al., 2019; McGranahan et al., 2016; Wang et al., 2019; Wang and Loreau, 2016, 2014; Wilcox et al., 2017; Zhang et al., 2019). Thus, there is an urgent need to understand temporal stability and the factors maintaining it at spatial scales covering larger areas (Gonzalez et al., 2020; Isbell et al., 2017; Wang et al., 2019).

**Figure 1.**
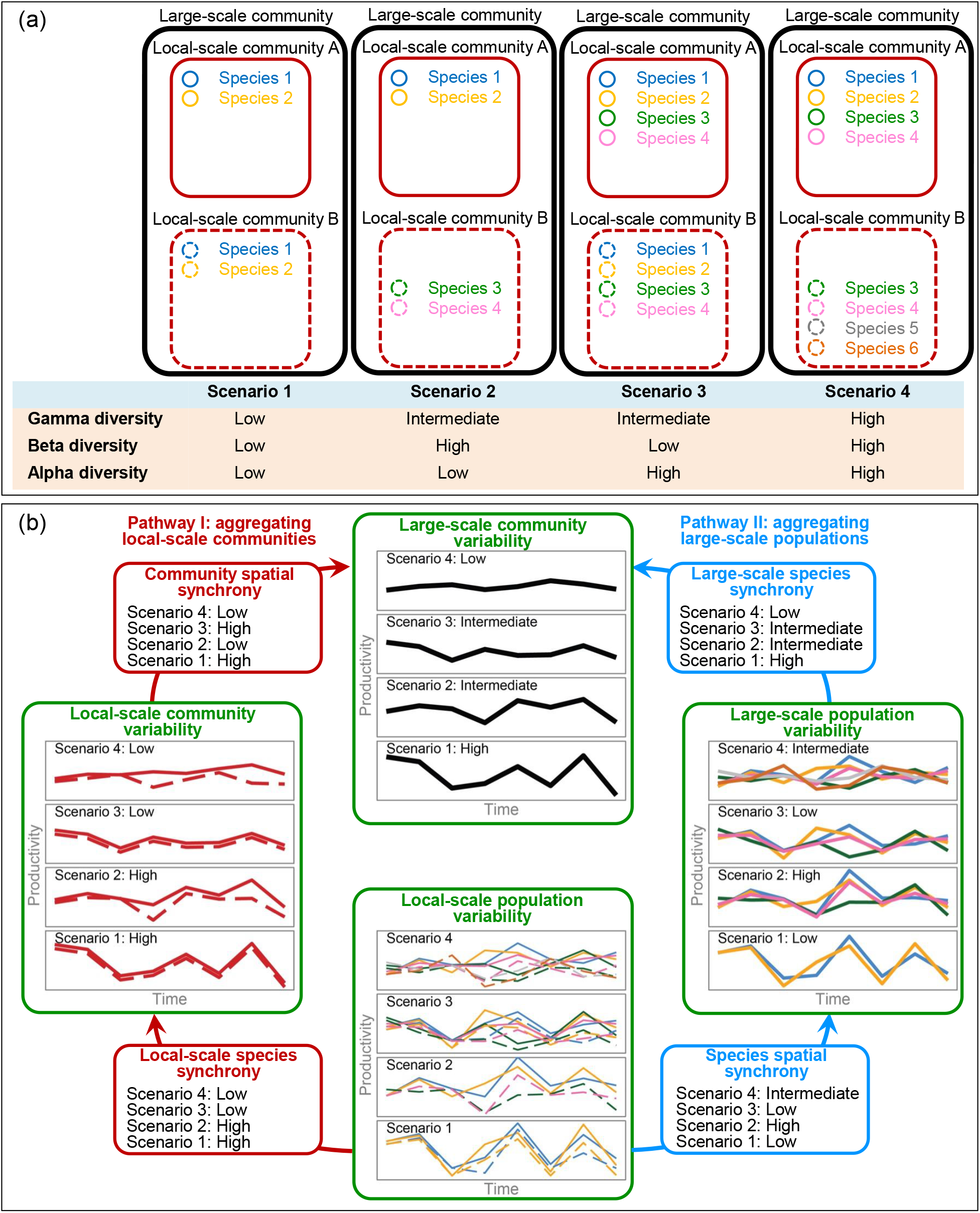
Diagrams showing large-scale communities with different scenarios of species diversity across spatial scales (a) and the large-scale community variability (estimated with coefficients of variation, CV, inverse of temporal stability) upscaled from local-scale population variability via local-scale communities (Pathway I, red arrows on the left side) and large-scale populations (Pathway II, blue arrows on the right side) under different scenarios (b; for terminology see Table 1). The subfigure (b) also shows theoretically proposed degrees of variabilities and synchronies (inverse of asynchrony) across ecological hierarchical levels under different scenarios (Thibaut and Connolly, 2013; Wang et al., 2019, 2020; Wang and Loreau, 2016, 2014). Mathematical derivations can be found in Supplementary file 2.

**Table 1.** Notation summary for climatic factors, species diversity indices, (temporal) coefficients of variation (CVs, inverse of temporal stabilities) and synchronies (inverse of asynchronies) across spatial scales and hierarchical levels of ecological organization. Details for estimating dominant-species components of CVs and synchronies can be found in Supplementary file 2.

| Symbol | Description |
| --- | --- |
| **Climatic factors** |  |
| $MGT$ | Cross-site averaged temporal mean growing-season temperature |
| $MGP$ | Cross-site averaged temporal mean growing-season precipitation |
| $CV_T^L$ | Local-scale temporal CV of (growing-season) temperature |
| $CV_P^L$ | Local-scale temporal CV of (growing-season) precipitation |
| $\phi_T^{L \rightarrow A}$ | Spatial synchronous dynamic of (growing-season) temperature |
| $\phi_P^{L \rightarrow A}$ | Spatial synchronous dynamic of (growing-season) precipitation |
| $CV_T^A$ | Large-scale temporal CV of (growing-season) temperature |
| $CV_P^A$ | Large-scale temporal CV of (growing-season) precipitation |
| **Biodiversity indices** |  |
| $N^\alpha$ or $N_d^\alpha$ | Alpha species richness or alpha dominant species richness |
| $N^\beta$ or $N_d^\beta$ | Beta species richness or beta dominant species richness |
| $N^\gamma$ or $N_d^\gamma$ | Gamma species richness or gamma dominant species richness |
| $D^\alpha$ or $D_d^\alpha$ | Alpha effective species richness or alpha dominant effective species richness |
| $D^\beta$ or $D_d^\beta$ | Beta effective species richness or beta dominant effective species richness |
| $D^\gamma$ or $D_d^\gamma$ | Gamma effective species richness or gamma dominant effective species richness |
| **Stability and synchrony** |  |
| $CV^{P,L}$ or $CV_d^{P,L}$ | **Local-scale population CV** or dominant-species local-scale population CV, defined as the weighted average local-scale population temporal CV estimated with all species or only dominant species within local-scale communities |
| $\phi^{P \rightarrow C,L}$ or $\phi_d^{P \rightarrow C,L}$ | **Local-scale species synchrony** or local-scale dominant species synchrony, defined as the weighted average synchronous dynamics among local-scale populations of all species or only dominant species within local-scale communities |
| $CV^{C,L}$ or $CV_d^{C,L}$ | **Local-scale community CV** or dominant-species local-scale community CV, defined as the weighted average community temporal CV estimated with all species or only dominant species |
| $\phi^{C,L \rightarrow A}$ or $\phi_d^{C,L \rightarrow A}$ | **Community spatial synchrony** or dominant-species community spatial synchrony, defined as the all-species or dominant-species estimates of weighted average spatial synchronous dynamics among local-scale communities |
| $\phi^{P,L \rightarrow A}$ or $\phi_d^{P,L \rightarrow A}$ | **Species spatial synchrony** or dominant species spatial synchrony, defined as the all-species and dominant-species estimates of weighted average spatial synchronous dynamics among local-scale populations |
| $CV^{P,A}$ or $CV_d^{P,A}$ | **Large-scale population CV** or dominant-species large-scale population CV, defined as the all-species and dominant-species estimates of weighted average population temporal CV at larger spatial scales |
| $\phi^{P \rightarrow C,A}$ or $\phi_d^{P \rightarrow C,A}$ | **Large-scale species synchrony** or large-scale dominant species synchrony, defined as the all-species and dominant-species estimates of weighted average synchronous dynamics among large-scale populations |
| $CV^{C,A}$ or $CV_{d,C}^{C,A}$<br>and $CV_{d,P}^{C,A}$ | **Large-scale community CV** or its dominant-species counterparts estimated via aggregating local-scale communities ($CV_{d,C}^{C,A} = \phi_d^{C,L \rightarrow A} \times CV_d^{C,L}$) or organizing large-scale populations ($CV_{d,P}^{C,A} = \phi_d^{P \rightarrow C,A} \times CV_d^{P,A}$) |

Recent theoretical work facilitates investigations of ecosystem temporal stability at a larger spatial scale by relating it to its hierarchical components along two alternative pathways I or II (Wang et al., 2019) (Figure 1b; see Table 1 for definition of terms used in this study). Along pathway I, in a first step asynchronous dynamics among different species populations due to their dissimilar responses (species insurance effect) (Tilman et al., 2014; Yachi and Loreau, 1999) stabilize communities at local scale. In a second step, spatial asynchronous dynamics among local communities due to heterogeneities in habitat and species composition (spatial insurance effect of communities) stabilize communities at a larger spatial scale (Wang and Loreau, 2016, 2014) (Figure 1b). Along pathway II, in a first step asynchronous dynamics among spatially separated local populations of each species, due to environmental heterogeneity (spatial insurance effect of populations) (Wang and Loreau, 2016, 2014), stabilize populations at a larger spatial scale. In a second step, asynchronous dynamics among large-scale populations of different species (species insurance effect) (Tilman et al., 2014; Yachi and Loreau, 1999) stabilize the large-scale community (Figure 1b). In the perhaps less likely case that populations and local communities respond synchronously to environmental fluctuations or environmental heterogeneity, the large-scale communities may be destabilized along the two alternative pathways. For example, a recent study showed that due to the strong driving effects of precipitation on biomass production of key species, its interannual variation forced synchronous dynamics of different species, destabilizing local communities (Wang et al., 2020).

Species diversity has been theoretically proposed to stabilize ecosystems at different ecological hierarchies, because species-rich communities are more likely to include species that have different responses to different environmental conditions across time and space, producing stable communities via species asynchrony (Thibaut and Connolly, 2013; Tilman et al., 2014; Wang and Loreau, 2016, 2014; Wang et al., 2020) (Figure 1b). In natural ecosystems, the role of species diversity in affecting temporal stability across different ecological hierarchies is still unclear. Theoretical and experimental studies propose stabilizing effects of (alpha) diversity within local communities (Hautier et al., 2015, 2014; Hector et al., 2010; Tilman et al., 2014, 2006) (Figure 1b). However, these studies usually consider systems in which species abundance distributions are relatively even, at least at the beginning, whereas natural communities are often characterized by highly uneven abundance distributions and dominated by the dynamics of a few abundant species (Thibaut and Connolly, 2013; Wang et al., 2019). In this case, the predicted local-scale diversity–stability relationship may be more difficult to be detected or it may be necessary to focus on the dynamic behavior of dominant species instead of species richness giving equal weight to all species (Wang et al., 2020; Xu et al., 2015; Yang et al., 2012). For example, our recent investigation on mechanisms maintaining temporal stability of local community biomass production in natural grasslands showed strong effects of dominant-species population dynamics instead of species richness (Wang et al., 2020). Furthermore, theoretical studies also propose that the heterogeneity in species compositions between spatially separated local communities (beta diversity) can increase asynchronous dynamics among them, resulting in stabilized communities at a larger spatial scale (Wang et al., 2019; Wang and Loreau, 2016) (Figure 1b). Currently, empirical evidence for such an effect is mixed as it was detected in some (Hautier et al., 2020; Liang et al., 2021; Wang et al., 2019) but not in other recent studies (Wilcox et al., 2017; Zhang et al., 2019). These studies looked at rather small spatial scales with potentially low beta diversity or even the same dominant species occurring among all local communities, making it difficult to detect a stabilizing effect of beta diversity. This further questions the usefulness of insights gained from studies across small spatial scales, even if they consider multiple local communities, for guiding regional management. Because different species may be dominant in different local communities in a larger spatial area, asynchrony among these local communities may contribute to temporal stability at a larger spatial scale (Wang et al., 2019; Wang and Loreau, 2016, 2014), a kind of spatial insurance (Isbell et al., 2018).

To investigate the temporal stability of biomass production (short “productivity”) at larger spatial scales, we established a region-scale observation network in Inner Mongolian grassland in China across an area of >166’894 km^2^ and monitored the yearly dynamics of productivity over five consecutive years (Figure 2a). The Inner Mongolian grassland represents a typical part of the Eurasian grassland biome and is crucial in providing biological products and services to human societies living there (Fang et al., 2015; Kang et al., 2007). In this region, plant community productivity and species richness and composition are driven by climatic factors, i.e. temperature and precipitation (Bai et al., 2004; Hu et al., 2018; Ma et al., 2010; Wang et al., 2020; Xu et al., 2015). These have changed considerably during the past decades (Huang et al., 2015; Piao et al., 2010) with largely unknown ecological consequences, especially at large spatial scales. To facilitate the large-scale temporal stability investigation, we employed a simulated landscape method (Hautier et al., 2018; van der Plas et al., 2016) to construct large-scale communities consisting of two local communities (two observed sites) separated by 17–987 km (Figure 2a). Briefly, each large-scale community was constructed by randomly choosing two local communities without replacement to ensure the constructed large-scale communities were independent between each other (see Figure 2b for a simplified 7-site case and Materials and Methods for details). Based on the above logical framework, we investigated how asynchronous dynamics among local or large-scale populations, especially those of dominant species (see Supplementary file 1–2 for details of dominant species and dominant-species measures), and among local communities contributed to the temporal stability of large-scale communities in the study region (see Supplementary file 3 for impacts of spatial distance). We used measures of synchrony and the coefficient of variation, CV, as “negative” proxies of asynchrony and temporal stability, respectively, and tested how these were affected by temporal variation in precipitation. We also tested whether species diversity could drive temporal stabilities at different spatial scales.

**Figure 2.**
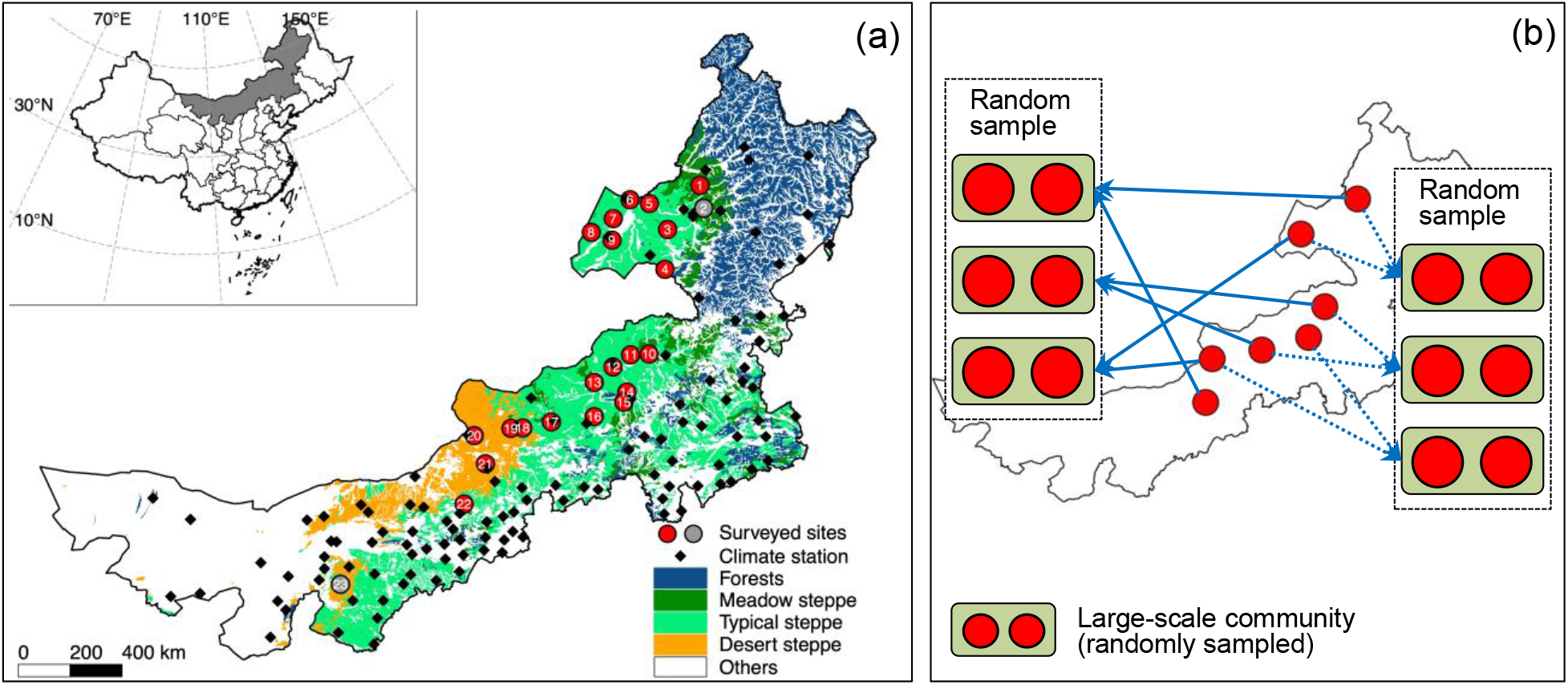
Geographical distribution of surveyed sites with site numbers (a) and a simplified case (7-site) for illustrating construction of large-scale communities aggregating two local-scale communities (b). In subfigure (a), red circles represent sites included in constructing large-scale communities (two sites, 2 and 23, with grey circles were excluded because they were monitored for only three years). The subfigure (b) shows a simplified case illustrating the construction of large-scale communities with a random resampling method without repeatedly using the same site to ensure constructed large-scale communities are independent between each other (see Materials and Methods for details).

## Results

We found that the large-scale community CV was positively associated with either all-species (Figure 3a–b, Figure 4a) or dominant-species measures (Figure 4b, see Supplementary file 2 for details of dominant-species measures) of local-scale community CV and community spatial synchrony in regression analyses and final SEMs based on the upscaling pathway I of aggregating local communities (see Figure 4–source data 1–2 for details of SEMs). In addition, the local-scale community CV (Figure 3e–f, Figure 4a) and its dominant-species counterpart (Figure 4b) were positively related to the local-scale population CV and local-scale species synchrony of all and dominant species, respectively. Furthermore, for all-species measures, the CVs decreased from 0.76 at the local population to 0.38 at the local community level and further to 0.29 at the large-scale community level (Figure 4a). We found that, in this upscaling pathway I, the local-scale species synchrony (mean = 0.49) was lower than the community spatial synchrony (mean = 0.78) (Figure 4a).

**Figure 3.**
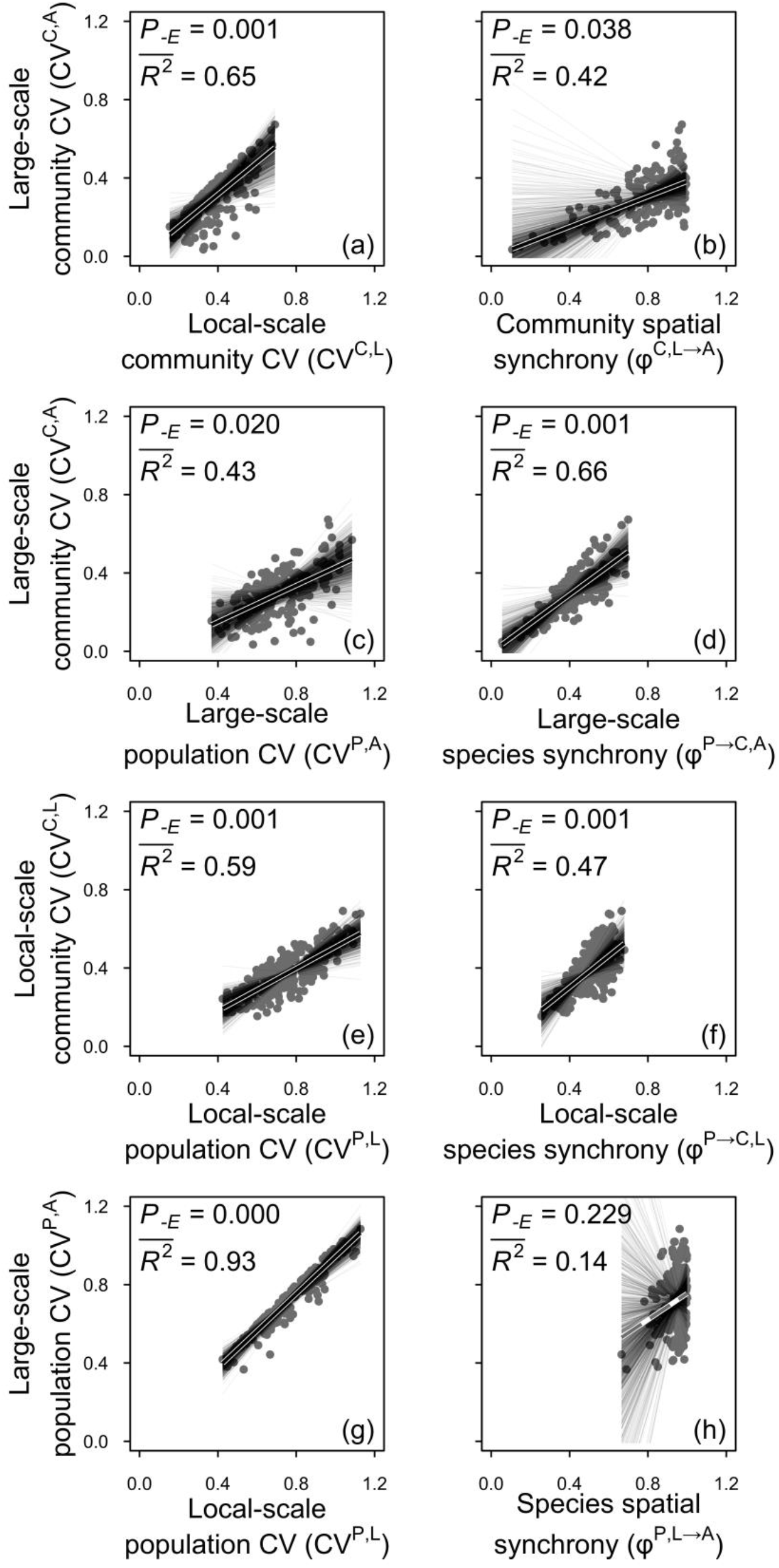
The large-scale community (a–d), local-scale community (e–f) and large-scale population (g–h) coefficients of variation (CVs, inverse of temporal stability) in relation to their hierarchical components. Solid black lines represent significant (*P* < 0.05) and marginally significant (*P* < 0.10) relationships and dashed grey lines represent non-significant (*P* > 0.10) relationships (see Materials and Methods for details and Table 1 for terminology). Dataset, code and relevant results can also be found in Figshare at https://doi.org/10.6084/m9.figshare.16903309.

**Figure 4.**
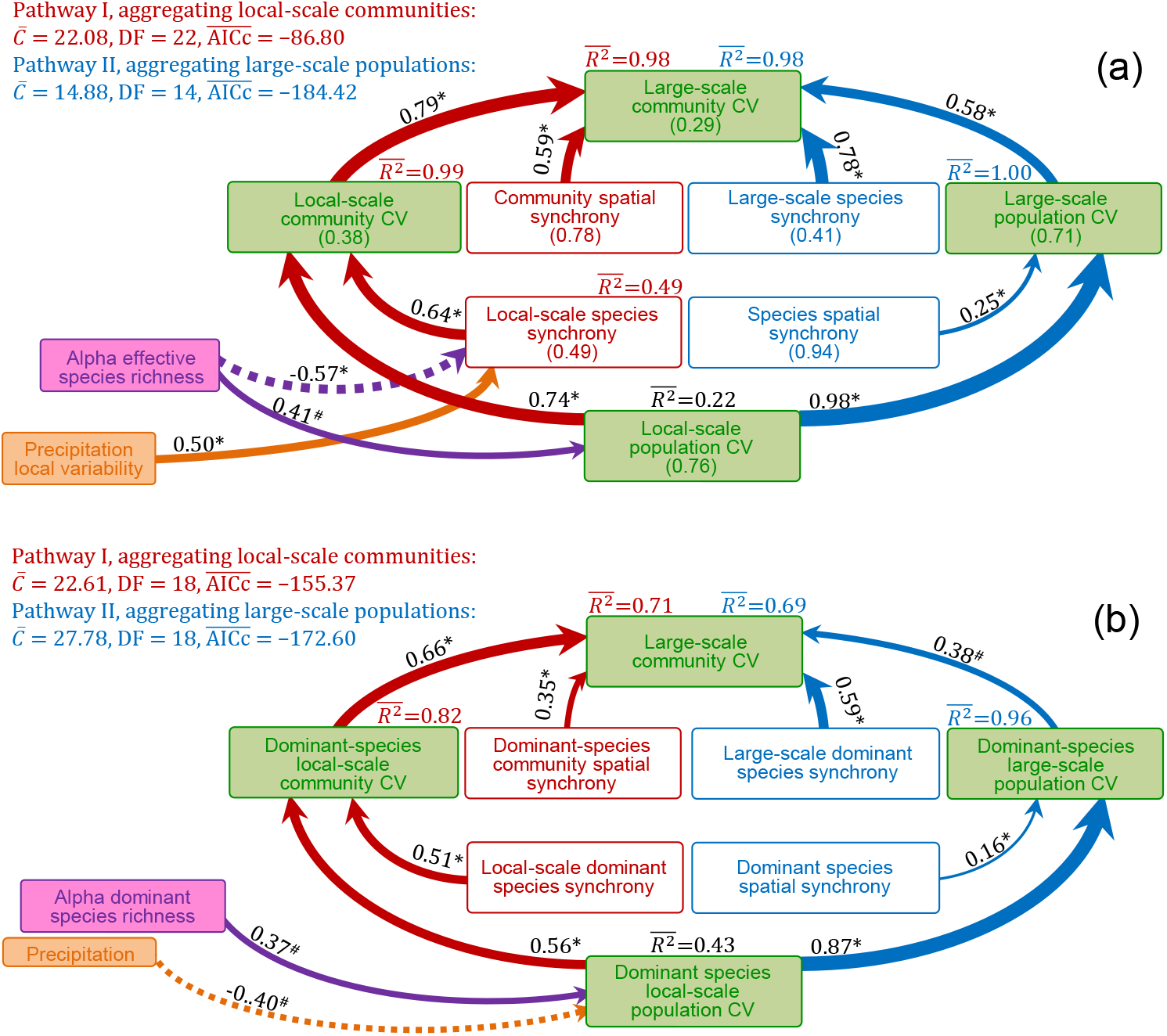
Diagrams of final structural equation models (SEMs) relating the large-scale community coefficient of variation (CV, inverse of temporal stability) to all-species (a) and dominant-species (b) measures of CVs and synchronies (inverse of asynchronies) at lower hierarchical levels of ecological organization and to species diversity indices and climatic factors. These diagrams combined pathways of local-scale population via local-scale community (upscaling pathway I on the left side with red arrows) and via large-scale population upscaling (upscaling pathway II on the right side with blue arrows) to the large-scale community (details of path analyses and initial and final SEMs that separately considering different upscaling pathways can be found in Figure 4–source data 1–2 and Figure 4–figure supplement 1–5). Solid and dashed arrows, respectively, represent examined positive and negative paths (see Figure 4–source data 1–2 for details). Arrows have also been scaled in relation to the strength of the relationship with numbers showing the mean values the standardized path coefficients. In addition, for all-species measures (a), mean values of CVs and synchronies are shown in brackets. The significance level of each path is indicated by * when *P* < 0.05 or # when *P* < 0.10 (see Materials and Methods for details). Dataset, code and relevant results can also be found in Figshare https://doi.org/10.6084/m9.figshare.16903309.

For the upscaling pathway II of aggregating large-scale populations, our final SEMs using all-species (Figure 3c–d, Figure 4a) and dominant-species measures (Figure 4b, see Supplementary file 2 for details of dominant-species measures) showed that the large-scale community CV was positively associated with the large-scale population CV and the large-scale species synchrony (see Figure 4–source data 1–2 for details of SEMs). In addition, although linear regression for all-species measures showed that the large-scale population CV was not related to species spatial synchrony (Figure 3h), this path was supported by the final SEM (Figure 4a). Furthermore, the CVs declined from 0.76 at the local-scale population level to 0.71 at the large-scale population level, and further to 0.29 at the large-scale community level (Figure 4a). In this upscaling pathway II, the large-scale species synchrony (mean = 0.41) was much lower than the species spatial synchrony (mean = 0.94) (Figure 4a).

We found that species diversity indices had almost no impacts on CVs and synchronies across ecological organization levels with few exceptions at the local scale, such as the impacts of local community diversity (i.e. alpha diversity) on local-scale species synchrony and local-scale population CV (see Materials and Methods for calculating species diversity indices across scales and Figure 4–source data 1–2 for details of SEMs). Specifically, gamma, beta and alpha diversity indices had no impacts on large-scale community CV, community spatial synchrony and local-scale community CV, respectively, when using either all-species (Figure 5a, 5d–5e, Figure 6) or dominant-species measures (Figure 4–figure supplement 1b). In addition, when using all-species measures, alpha species diversity negatively influenced local-scale species synchrony but positively influenced local-scale population CV (Figure 4a, Figure 5f–5g, Figure 6b). When using dominant-species measures, only the alpha species richness had a positive impact on local-scale population CV (Figure 4b). Moreover, gamma diversity indices had no influences on large-scale species synchrony when using either all-species (Figure 5c and 6, Supplementary file 4) or dominant-species measures (Figure 4–figure supplement 1b). In addition, correlation and regression analyses showed that large-scale population CV was positively associated with gamma diversity when using all-species measures (Figure 5b) and positively associated with gamma species richness when using dominant-species measures (Figure 4–figure supplement 1). However, these paths were not supported by the final SEMs (Figure 4a–4b). Besides, our SEMs (Figure 6) and general linear models (Supplementary file 4) further exploring the impacts of species diversity indices across ecological hierarchies showed no impacts on the CVs of local community, large-scale population and large-scale community. We further found that dominant species as a group had strong impacts on CVs and synchronies with mean explanatory power 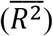 generally > 0.52 (Supplementary file 5), expect for the dominant species spatial synchrony (*P_−E_* = 0.17, 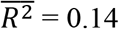, Supplementary file 5–Figure 1h).

**Figure 5.**
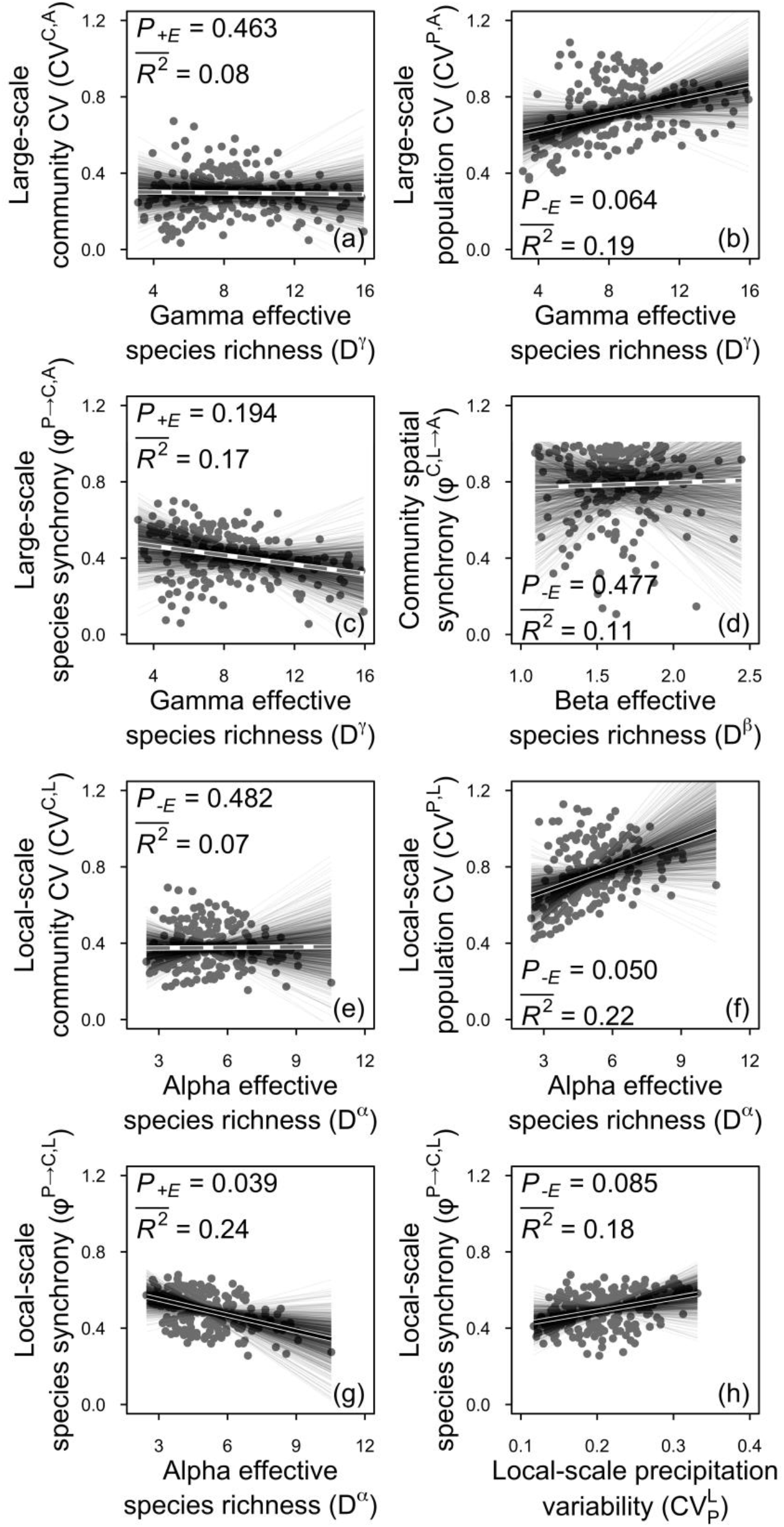
Coefficients of variation (CVs, inverse of temporal stability) and synchronies (inverse of asynchrony) across spatial scales in relation to species diversity (effective species richness, a–g) and local-scale species synchrony in relation to local-scale precipitation variability (h). Solid black lines represent significant (*P* < 0.05) and marginally significant (*P* < 0.10) relationships and dashed grey lines represent non-significant (*P* > 0.10) relationships (see Materials and Methods for details). Dataset, code and relevant results can also be found in Figshare https://doi.org/10.6084/m9.figshare.16903309.

**Figure 6.**
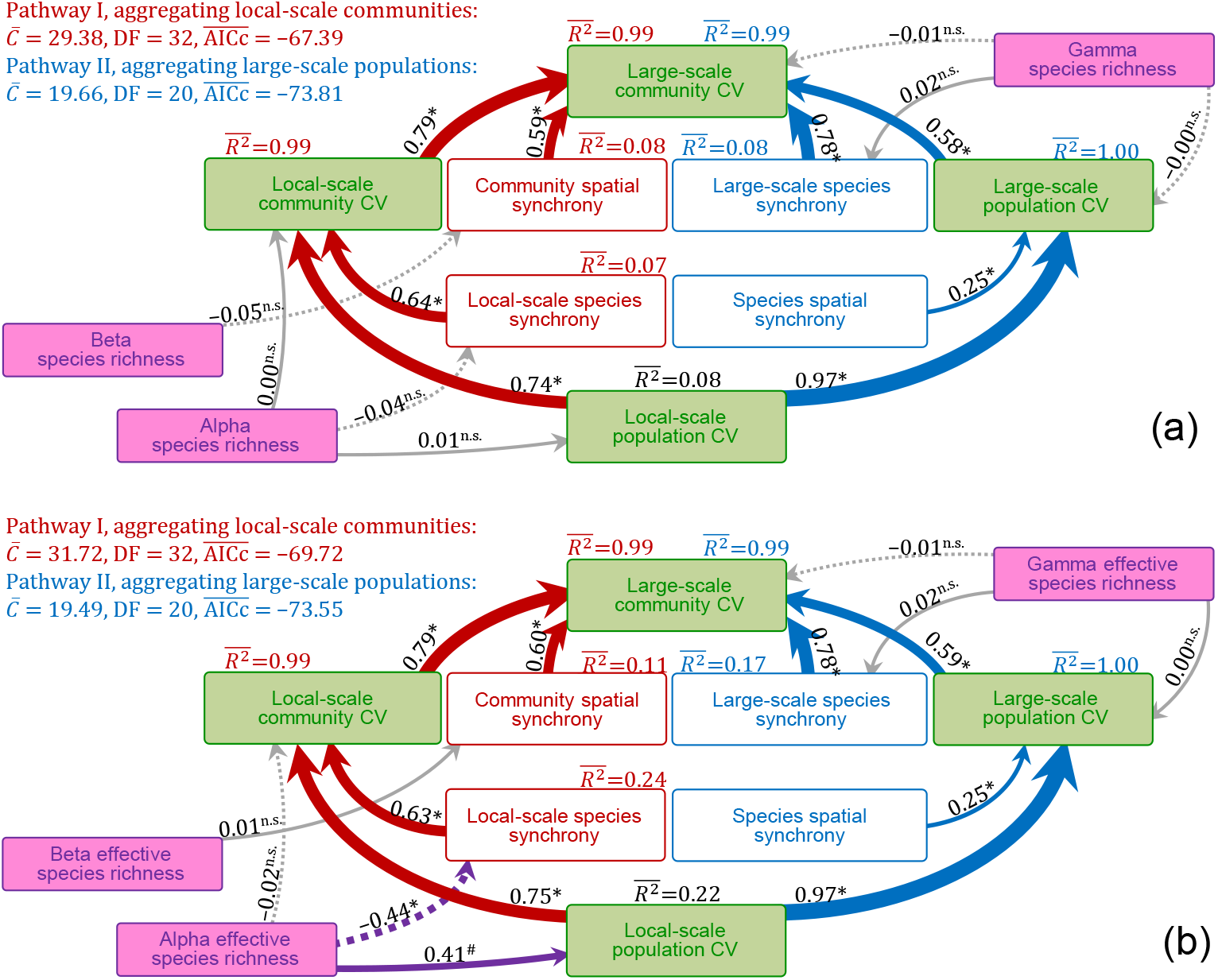
Diagrams of structural equation models (SEMs) examining theoretically proposed impacts of species diversity (species richness, a, and effective species richness, b) on the large-scale community coefficient of variation (CV, inverse of temporal stability) and its hierarchical components. These diagrams combined local-scale population via local-scale community (upscaling pathway I on the left side with red arrows) and via large-scale population (upscaling pathway II on the right side with blue arrows) upscaling to the large-scale community (details of separately considering different upscaling pathways can be found in Figure 6–source data 1 and Figure 6–figure supplement 1). Colored and grey arrows represent significant (or marginally significant) and non-significant paths, respectively. Solid and dashed arrows, respectively, represent examined positive and negative paths (Figure 6–source data 1). Arrows have also been scaled in relation to the strength of the relationship with numbers showing the mean values the standardized path coefficients. The significance level of each path is indicated by * when *P* < 0.05, # when *P* < 0.10 or n.s. (non-significant) when *P* > 0.10 (see Materials and Methods for details). Dataset, code and relevant results can also be found in Figshare https://doi.org/10.6084/m9.figshare.16903309.

We also found that large-scale CVs and spatial synchronies of growing-season temperature and precipitation had no impacts on large-scale community CV and its hierarchical components (Figure 4–figure supplement 1). However, within local communities, local-scale species synchrony increased with local-scale precipitation variability (Figure 4a, Figure 5h), whereas the local-scale population CV of dominant species was reduced by larger mean values of precipitation (Figure 4b, see Figure 4–source data 1–2 for details of SEMs).

## Discussion

Based on a region-scale survey over 5 years in Inner Mongolian grassland, we investigated temporal stabilities (inverse of CVs) and asynchronies (inverse of synchronies) across spatial scales, and analyzed influences of species diversity, abundant species and climatic factors on them. We found that temporal stabilities at large spatial scale, i.e. large-scale community temporal stability, was related to that of and asynchronous dynamics among units at small scale, i.e. local-scale community temporal stability and community spatial asynchrony. However, stabilities and asynchronies were only impacted by species diversity at local scale but were driven by dominant species at local and large scales. Furthermore, decreasing mean and increasing interannual fluctuation of precipitation could, respectively, destabilize dominant species and synchronize population dynamics within local communities, impairing stability at large scales. These results indicate that reduced amounts and increased variation of precipitation (Huang et al., 2015; Piao et al., 2010) are key climatic changes threatening the sustainable delivery of biological products and services to human well-being in this region.

### Stability across ecological hierarchies

We investigated stabilities across ecological hierarchies with two alternative upscaling pathways (Wang et al., 2019) and both of them showed gradually increasing temporal stability from low to high organization levels due to species insurance effects and spatial insurance effects of populations and communities, caused by asynchronous dynamics among species and localities (Figure 4a). These patterns are consistent with recent studies carried out at single sites constructing multiple adjacent plots within meta-communities in grassland ecosystems (Hautier et al., 2020; McGranahan et al., 2016; Wang et al., 2019; Wilcox et al., 2017; Zhang et al., 2019) and at the regional scale in marine ecosystems (Lamy et al., 2019; Thorson et al., 2018), as well as recent theoretically proposed positive invariability–area relationships (Isbell et al., 2018; Wang et al., 2017). These results suggest that, at large spatial scales, spatial heterogeneity is important in maintaining stability; losing this heterogeneity (Fahrig et al., 2011; Gámez-Virués et al., 2015) can impair stability.

We found that the species insurance effect caused by among-species dissimilar responses (Tilman et al., 2014; Yachi and Loreau, 1999) was stronger in maintaining temporal stability at large spatial scales than the spatial insurance effects of populations and communities, despite the large spatial extent and thus expected large spatial heterogeneity of our study region (Figure 2a). This result is consistent with a recent investigation in marine plant communities (Lamy et al., 2019) but different from that in fish communities (Thorson et al., 2018). In our study, the region-wide synchronous variations in precipitation (mean = 0.86, ranged from 0.62 to 1.00) (Supplementary file 3–Figure 1b) potentially decreased the spatial heterogeneity and increased the relative importance of among-species dissimilarity and the species insurance effect. The more mobile fish populations and communities (Thorson et al., 2018) may be strongly attracted by certain environmental conditions, causing largely different spatial population patterns across years, strengthening the spatial insurance effects of populations and communities. In plant communities, the strong species insurance effect suggests that the large-scale community stability at least partly reflects the stability of local communities, which are prevalent in previous studies (Ma et al., 2017; Tilman et al., 2006; Xu et al., 2015; Yang et al., 2012). However, the large-scale community stability does not so much reflect local population and large-scale population stabilities. The species insurance effect has also been shown to regulate ecosystem resistance and resilience to extreme climate evens, e.g. drought (Xu et al., 2014). Our results indicate that insights on local-scale resistance and resilience (Isbell et al., 2015) can also potentially reflect these characteristics of larger spatial scales.

### Influence of species diversity, dominant species and precipitation on ecosystem stability

We only detected stabilizing impacts of species diversity at local scale (Figure 4, Figure 5f–5g, Figure 6). The negatively impacted local-scale population temporal stability by alpha diversity is in line with theoretical and experimental biodiversity studies (Lehman and Tilman, 2000; Tilman, 1999; Tilman et al., 2014, 2006), proposing that competition between coexisting species for resources in species-rich communities leads to low population stability. In addition, the detected positive association between local-scale species asynchrony and alpha diversity potentially results from the higher probability of species-rich communities to contain species that are different in responding to environmental fluctuations (Tilman et al., 2014; Yachi and Loreau, 1999).

Previous studies reported mixed impacts of species diversity on stabilities and asynchronies at scales beyond the local. Some studies found significant influences (Hautier et al., 2020; Liang et al., 2021; Wang et al., 2019) and others found none (Wilcox et al., 2017; Zhang et al., 2019). It has been argued recently (Hautier et al., 2020) that investigations within a single site (Zhang et al., 2019) or multiple sites with non-standardized experimental protocol (Wilcox et al., 2017) may mask stabilizing effects of species diversity at large spatial scales. The current study used a multi-site dataset with a standardized survey protocol and found no impacts of species diversity at scales beyond local (Figure 4, Figure 6) but strong driving effects of dominant species at local and large scales (Figure 4b, Supplementary file 5). The highly uneven distribution of species abundances could have been responsible for this pattern (Wang et al., 2020), as under the uneven distribution the contribution of the most diverse part of the community to stabilities and asynchronies was limited by low abundance (Thibaut and Connolly, 2013; Wang et al., 2019). Considering that many natural ecosystems are characterized by high unevenness (Dee et al., 2019; Jiang et al., 2009; Smith and Knapp, 2003), the reported strong influences of abundant species and weak influences of all-species diversity on stabilities and asynchronies may be quite common in the real world. More importantly, the current study also provides a tool to quantify impacts of abundant species, or even a certain species or a certain functional group, on stabilities and asynchronies at different ecological hierarchies.

The strong influence of precipitation on productivities of different species (Zhang et al., 2017) may also weaken the (spatial) insurance effect of species diversity (i.e. local-scale species asynchrony). Under such circumstances, fluctuation in precipitation forces similar responses in different species, decreasing the dissimilarity and thus asynchrony among species. This speculation is supported by the low local-scale species asynchrony under high precipitation fluctuation (Figure 4a, Figure 5h). Furthermore, we also found decreased dominant-species local-scale population temporal stability under low precipitation amount (Figure 4b), potentially owing to the decreasing mean-to-standard deviation ratio caused by the dominant-species biomass production being more steeply related to precipitation amount than to its standard deviation (Wang et al., 2020). The study region has been experiencing a pronounced decrease in precipitation and an increase in its variability during the past decades (Huang et al., 2015; Piao et al., 2010; Tao et al., 2015). Our results indicate that these changes in precipitation regimes may present a key threat to the sustainable provision of biological products and services to human well-being in the region.

## Materials and methods

### Study region and plant community survey

The Inner Mongolian temperate grassland has a continental monsoon climate with a short and cool growing season (from May to October, averaged temperature 12.9–18.4 °C across sites during the studied period from 2012–2016), concentrating ~90% of the annual precipitation (averaged precipitation 186.2–398.0 mm across sites from 2012–2016) (Wang et al., 2020). This region has three main vegetation types: meadow steppe (dominated by perennial grasses and forbs e.g. *Stipa baicalensis*, *Leymus chinensis* and *Convolvulus ammannii*), typical steppe (dominated by perennial grasses e.g. *Stipa grandis*, *Leymus chinensis* and *Stipa krylovii*) and desert steppe (dominated by perennial grasses and forbs e.g. *Stipa caucasica* and *Allium polyrhizum*) (Figure 2a). In this area, grazing and mowing are the most widely practiced land-use regimes with increasing intensities during the last decades (Fang et al., 2015; Wu et al., 2015).

We established a 5-year (2012–2016) region-scale survey over this area, including 23 individual sites (latitudes 39.34–49.96 °N, longitudes 107.56–120.12 °E) covering all three grassland types (Figure 2a) (Wang et al., 2020). The sample plots of each site were randomly selected, excluding heavy anthropogenic disturbances (e.g. grazing and mowing). The plant communities were surveyed between late July and early August in each year in the following way. At each site, we marked three 1 m × 1 m quadrats along the diagonal of a 10 m × 10 m plot, harvested all living plant tissues and sorted them to species, and then oven-dried and weighed the harvested material to obtain aboveground biomass and species richness (for details see (Wang et al., 2020)).

### Construction of large-scale communities

We constructed large-scale communities consisting of two local communities. We excluded 2 sites with only 3-year available data as their 2-year overlaps with others were too short for calculating a CV (Figure 2a), resulting in only 21 sites with available data of 4–5 years (2, 15 and 4 sites for meadow, typical and desert steppe, respectively) (Wang et al., 2020). The construction of large-scale communities was done with a simulated landscape method (Hautier et al., 2018; van der Plas et al., 2016). Specifically, the 21 local communities (sites) were randomly separated into 10 large-scale communities without replacement (2 local communities for each large-scale community with 1 remainder) to ensure that they were independent between each other (see Figure 2b for a simplified 7-site case). We repeated this random resampling process 1000 times, resulting in 1000 resampled sets, each containing 10 large-scale communities that were independent of each other.

### All-species and dominant-species diversity indices, CVs and synchronies across ecological hierarchies

We estimated two alternative species diversity indices across ecological hierarchies, species richness (*N*) and effective species richness (*D*). The alpha (*N^α^*) and gamma species richness (*N^γ^*) were defined as the total number of species at local and large scales and the beta species richness (*N^β^* = *N^γ^* / *N^α^*) was used to measure dissimilarity among localities. Specifically, the alpha (*N^α^*) and gamma (*N^γ^*) species richness were estimated as multiple-year mean (*N^α^*) and multiple-year pooled species number (*N^γ^*) of the two local communities. To account for highly uneven species abundances in the study region, we also used effective species richness, the antilog of Shannon-Wiener diversity (*D = e^H’^*), reflecting how many species with an even abundance distribution would produce the same Shannon-Wiener diversity as observed for the actual uneven community (Wang et al., 2020). The alpha (*D^α^*) and gamma (*D^γ^*) effective species richness thus represented the Shannon-Wiener diversity at local and large scales, with beta effective species richness (*D^β^* = *D^γ^* / *D^α^*) measuring its cross-locality dissimilarity (estimated with the same method used for species richness). These species diversity indices were estimated with all species or only dominant species, the latter defined as species whose biomass contributed > 5% to the total biomass of the large-scale community (Wang et al., 2020) (Supplementary file 1) over the 5 survey years (dominant-species measures designated with subscript *d*, such as 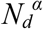 for the alpha dominant species richness).

Here, we illustrate a recent theoretical framework (Wang et al., 2019) upscaling local-scale population CV to large-scale community CV and use superscripts *P* and *C* to designate the quantities at population level and community level, superscripts *L* and *A* the quantities of localities (e.g. local communities) and an aggregation of multiple localities (e.g. large-scale communities). In addition, we used superscript *P→C* and *L→A* to designate upscaling processes of organizing populations and aggregating localities, respectively. This theoretical framework showed that local-scale population CVs (*CV^P,L^*) can be upscaled to the large-scale community CV (*CV^C,A^*) via either the dynamics of local communities (*CV^C,L^*) or via the dynamics of large-scale populations (*CV^P,A^*) (Figure 1b, see Table 1 for details of abbreviations). In the first upscaling pathway (pathway I), local populations were first organized into local communities and then local communities were aggregated into large-scale communities. In this process, the CV decreases from local population to local community level and further to the level of large-scale community. The degrees of these decreases are determined by synchronous dynamics among local populations of different species within local communities (local-scale species synchrony, *φ^P→C,L^*) and among spatially separated local communities (community spatial synchrony, *φ^C,L→A^*), respectively (Figure 1b). This is because synchronies take values between 0 (perfectly asynchronous) to 1 (perfectly synchronous), thus measuring the proportion of CVs upscaled to higher organization levels from local populations to local communities or local communities to large-scale communities (Wang et al., 2019). In the alternative upscaling pathway (pathway II), the local populations were first aggregated to large-scale populations and then the large-scale populations were organized into large-scale communities. In this process, the decreases of CVs are determined by synchronous dynamics among spatially separated local populations of same species (species spatial synchrony, *φ^P,L→A^*) and among large-scale populations of different species (large-scale species synchrony, *φ^P→C,A^*) (Figure 1b).

We extended this theoretical framework to separate CVs and synchronies into dominant and subdominant species groups (Supplementary file 2) and only investigated the contributions of the dominant-species group to CVs and synchronies of communities consisting only of dominant species because remaining species contributed very little to total biomass and reduced model fits and predictions (Thibaut and Connolly, 2013; Wang et al., 2019). Briefly, in the upscaling pathway of aggregating local communities (pathway I), the dominant-species local population 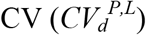 stepwise interacts with dominant-species measures of local-scale species synchrony 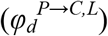 and community spatial synchronies 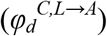 and upscales to the dominant-species local community 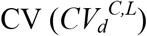 and large-scale community CV, respectively (Supplementary file 2A–2C). In the upscaling pathway of organizing large-scale populations (pathway II), the dominant-species local population CV stepwise interacts with dominant-species measures of species spatial synchrony 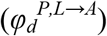 and large-scale species synchrony 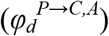 and upscales to the dominant-species large-scale population 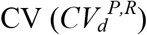 and large-scale community CV, respectively (Supplementary file 2D–2E). The two upscaling pathways can produce slightly different large-scale community CVs (Supplementary file 2F), which is why we use two abbreviations for the latter, i.e. 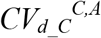 and 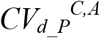 for upscaling pathways of aggregating local communities and organizing large-scale populations.

### Climatic data

Based on monthly climatic data collected from 119 climate stations and 2-km resolution digital elevation over this region, we calculated site-specific mean temperature and precipitation using the simple kriging method and spherical model of geostatistical analysis in ArcGIS software (Environmental Systems Research Institute Inc., Redlands, CA, USA). Because plants are more active during the growing season, only growing-season temperature (*MGT*), precipitation (*MGP*) and their CVs across spatial scales were used in the current study. Specifically, temperature (*MGT*) and precipitation (*MGP*) are cross-site averaged multi-year mean temperature and precipitation. In addition, CVs of *MGT* and *MGP* at the local (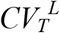 and 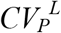) and large scales (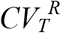 and 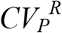), as well as their among-site synchronies (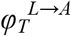 and 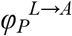) were estimated with the methods used for local-scale and large-scale community CVs and community spatial synchrony.

### Statistical analysis

We analyzed the influence of distance between spatially separated local communities (i.e. sites) within large-scale communities on spatial synchronies of *MGT* and *MGP*, large-scale community CV as well as all-species and dominant-species measures of large-scale population CV and species and community spatial synchronies with linear regressions. However, we did not further include distance between sites as explanatory term in statistical analyses. This is because spatial distance only influenced spatial synchronies of *MGT* and *MGP* (Supplementary file 3) and both of them were not included in initial path-analysis models (see below for details of path analysis and statistical significance).

We used correlation analyses, linear regressions and path analyses to investigate the large-scale community CV in relation to its hierarchical components, species diversity indices and climatic factors (Figure 4–figure supplement 1, see Figure 4–figure supplement 2 for details of scenario combining three local communities into a large-scale community). Specifically, we established initial path-analysis models separately considering different upscaling pathways and different species diversity indices, as well as the large-scale community CV and its hierarchical components estimated with all species or only dominant species (Figure 4–figure supplement 3–5). These initial models stayed as close as possible to paths proposed to be essential in correlation analyses and recent theoretical studies (Wang et al., 2019; Wang and Loreau, 2016, 2014) (Figure 4–source data 1–2, Figure 4–figure supplement 3–5). Then, structural equation models (SEMs, Figure 4–source data 1) and general linear models (Figure 4–source data 2) were used to analyze these proposed paths, to eliminate non-significant ones until containing only significant or marginally significant paths or reaching the lowest value of Akaike’s information criterion (for small sample size, *AICc*). Subsequently, SEMs were used to analyze the remaining paths in the final models (Figure 4–source data 1, Figure 4–figure supplement 5) (*piecewiseSEM* package (Lefcheck, 2016) of R 3.6.3 (R Core Team, 2013)). We note that the SEMs were only used to analyze the strengths of paths, rather than searching for best models *post hoc*. We did this even at the cost that overall model fits might have significantly deviated from a saturated model and used Shipley’s test of d-separation (Lefcheck, 2016; Shipley, 2013) besides Fisher’s *C* statistic (*C*) and *AICc* as an additional guide (Figure 4–source data 1).

Because species diversity indices were rarely included in initial path-analysis models (Figure 4–figure supplement 3–5), we used SEMs to further explore impacts of species diversity indices on the large-scale community CV and its hierarchical components based on theoretical predictions (Wang et al., 2019) (Figure 6–figure supplement 1, Figure 6–source data 1). In addition, we also used general linear models (Supplementary file 4) to further explore local-scale and large-scale community CVs in relation to species diversity indices, considering the influences of their hierarchical components based on theoretical predictions (Wang et al., 2019). Specifically, we investigated the large-scale community CV in relation to gamma and beta diversity indices, separately considering local community CVs and community spatial synchrony as well as population CV and species synchrony of local scale. In addition, within local communities, we also explored the community CV in relation to alpha diversity indices, population CV and species synchrony.

We used a randomized examination method to investigate the statistical significance of the above analyses. Specifically, considering the 10 independent large-scale communities per sampled set, all above statistical analyses were conducted within each set, resulting in 1000 statistics. These were then analyzed with the randomized examination method. Taking the correlation analysis as an example, we calculated the mean correlation coefficient 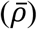 of the 1000 sets and considered it to be statistically significant or marginally significant if the proportion of *ρ* < 0 (*P_−ρ_*) (or *ρ* > 0, *P_+ρ_*) was lower than 0.05 or 0.10 when 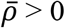 (or 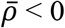), respectively. For linear regressions and SEMs, we also used the randomized examination method to analyze the statistical significances of the estimated coefficients and calculated the mean explanatory power 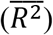 of them, as well as the mean Fisher’s *C* statistic 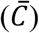 and the mean *AICc* 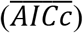 of SEMs.

## Acknowledgement

This study was supported by the National Nature Science Foundation of China (31960259, 31971434, 32160274, 31370454 and 31600385), the National Key Research and Development Program of China (2016YFC0500602), the Ministry of Science and Technology of China (2015BAC02B04) and the Natural Science Foundation of Inner Mongolia (2019MS03089, 2019MS03088 and 2015ZD05). S.W. was supported by the National Nature Science Foundation of China (31988102). B.S. was supported by the University Research Priority Program Global Change and Biodiversity of the University of Zurich. All authors declare no conflict of interest.

## Author contributions

YW, LZ, CL, BM, QZ, XN and WM designed the study and compiled the data. YW produced the results and wrote the first draft with SW, WM and BS. All authors contributed to the development of the manuscript.

## Data availability statement

The data that support the findings of this study are openly available in Figshare at https://doi.org/10.6084/m9.figshare.16903309.

## Supplementary Information

Additional supporting information may be found online in the Supporting Information section at the end of this article.

---

**Figure 4–figure supplement 1.**
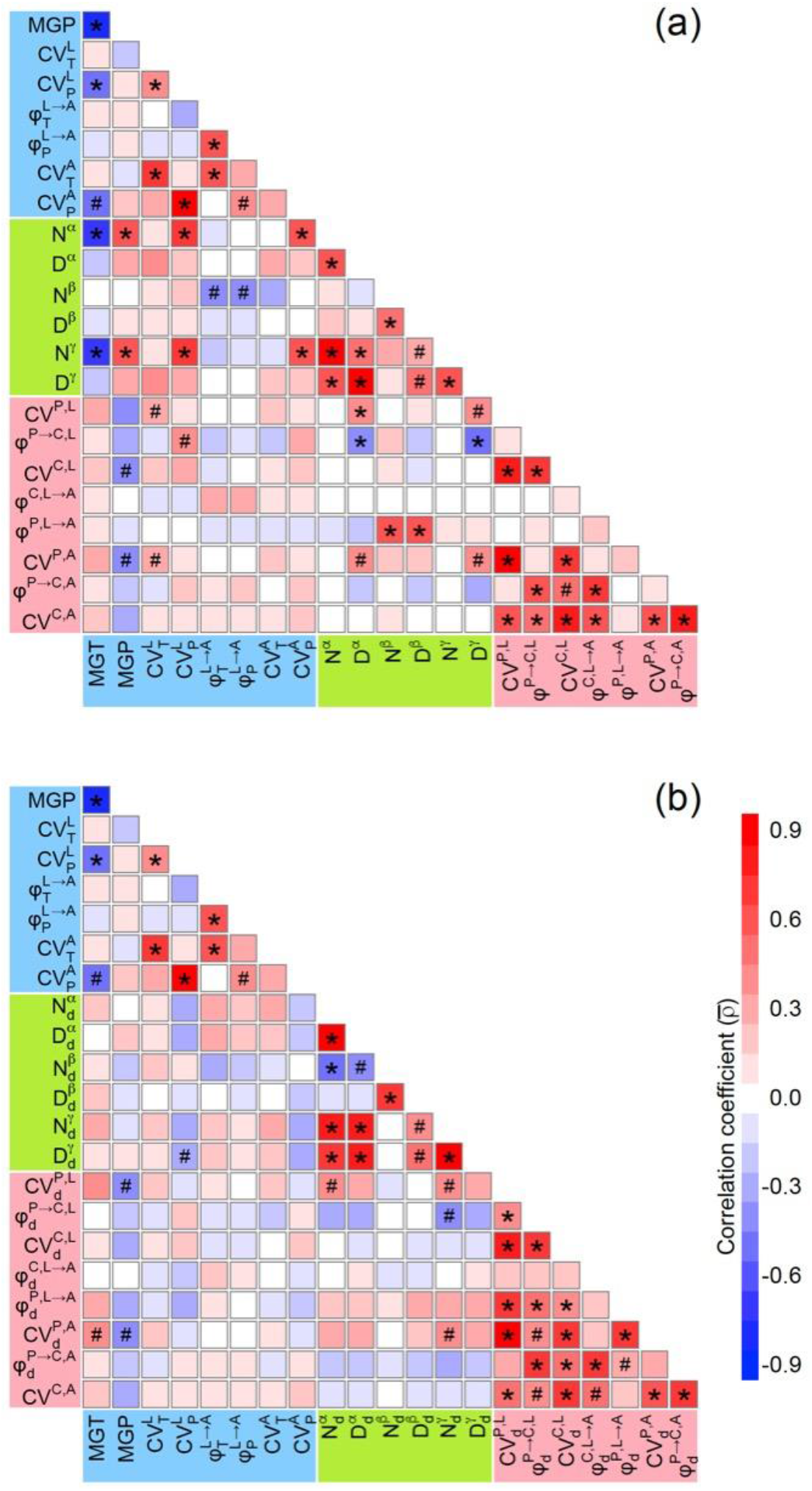
Correlation matrices for climatic factors, species diversity indices, coefficients of variation (CVs, inverse of temporal stabilities) and synchronies (inverse of asynchronies) estimated with all species (a) and only dominant species (b) by considering a 2-local-community scenario (see Figure 2b for a simplified case). Significant and marginally significant correlations are marked with * (*P* < 0.05) and # (*P* < 0.10), respectively (see Materials and Methods for details). Symbols and descriptions can be found in Table 1. Dataset, code and relevant results can also be found in Figshare https://doi.org/10.6084/m9.figshare.16903309.

**Figure 4–figure supplement 2.**
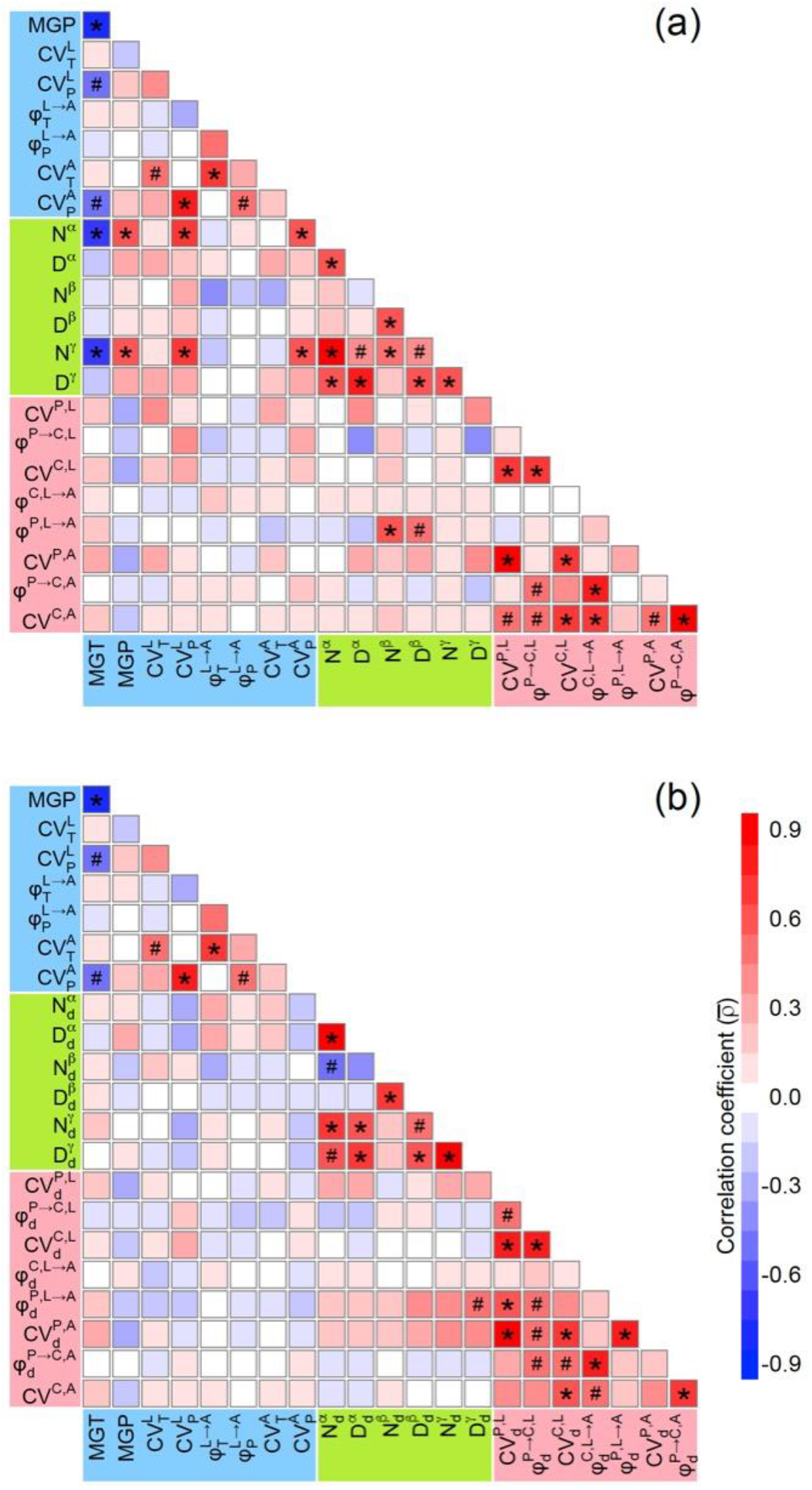
Correlation matrices for climatic factors, species diversity indices, coefficients of variation (CVs, inverse of temporal stabilities) and synchronies (inverse of asynchronies) estimated with all species (a) and only dominant species (b) by considering a 3-local-community scenario (similar sampling as in Figure 2b, but with seven groups of three sites per sample). Significant and marginally significant correlations are marked with * (*P* < 0.05) and # (*P* < 0.10), respectively (see Materials and Methods for details). Symbols and descriptions can be found in Table 1. Potentially owing to the small sample size (*n* = 7) of the 3-local-community scenario, many significant (or marginally significant) correlations showed in the 2-local-community scenario (*n* = 10, Figure 4–figure supplement 1) were non-significant. Thus, we did not further analyze the 3-local-community scenario. Dataset, code and relevant results can also be found in Figshare https://doi.org/10.6084/m9.figshare.16903309.

**Figure 4–figure supplement 3.**
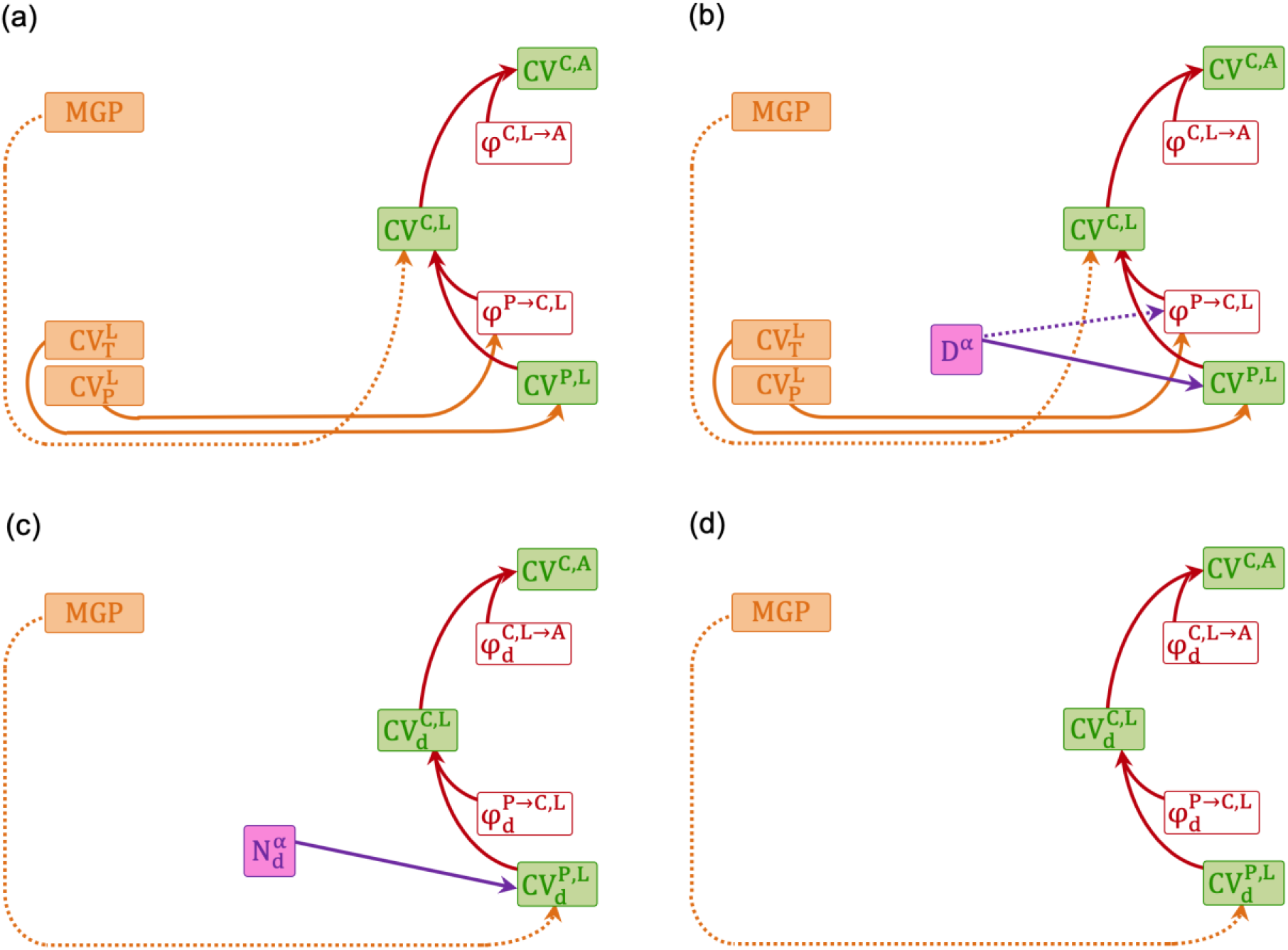
Initial structural equation models (SEMs) relating the large-scale community coefficient of variation (*CV^C,A^*, inverse of temporal stability) to its hierarchical components, species diversity indices and climatic factors using the upscaling pathway of aggregating local-scale communities (pathway I). These models considered CVs and synchronies (inverse of asynchronies) estimated with all species (a and b) or only dominant species (c and d). In addition, they also considered two alternative species diversity indices, species richness (*N*, a and c) and effective species richness (*D*, b and d). Solid and dashed arrows represent significant (or marginally significant) positive and negative correlation relationships, respectively (Figure 4– figure supplement 1). Because (b) includes all paths of (a) and (c) includes all paths of (d), only the models shown in (b, Figure 4–figure supplement 5a) and (c, Figure 4–figure supplement 5c) are further analyzed with SEMs (Figure 4–source data–1A–1B) and general linear models (Figure 4–source data–2A–2B). Symbols and descriptions can be found in Table 1.

**Figure 4–figure supplement 4.**
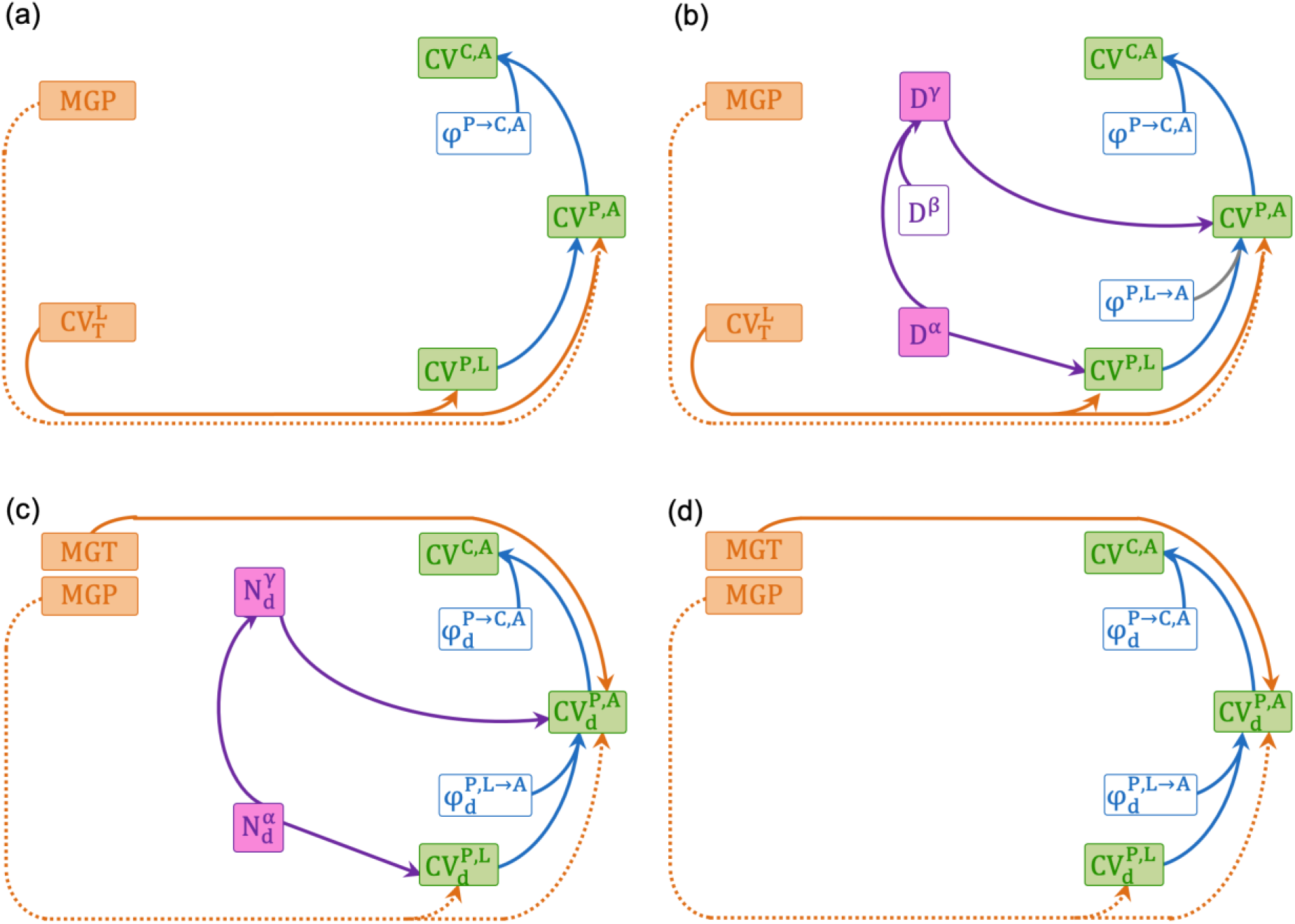
Initial structural equation models (SEMs) relating the large-scale community coefficient of variation (*CV^C,A^*, inverse of temporal stability) to its hierarchical components, species diversity indices and climatic factors using the upscaling pathway of organizing large-scale populations (pathway II). These models considered CVs and synchronies (inverse of asynchronies) estimated with all species (a and b) or only dominant species (c and d). In addition, they also consider two alternative species diversity indices, species richness (*N*, a and c) and effective species richness (*D*, b and d). Solid and dashed color arrows represent significant (or marginally significant) positive and negative correlation relationships, respectively (Figure 4– figure supplement 1). Grey solid arrow (large-scale population CV in relation to species spatial synchrony, b) represents non-significant positive correlation relationship, which is added in the initial structure equation model because it is theoretically proposed important (Wang et al., 2019). Because (b) includes all paths of (a) and (c) includes all paths of (d), only the models shown in (b, Figure 4–figure supplement 5e) and (c, Figure 4–figure supplement 5g) are further analyzed with SEMs (Figure 4–source data–1C–1D) and general linear models (Figure 4–source data–2C–2D). Symbols and descriptions can be found in Table 1.

**Figure 4–figure supplement 5.**
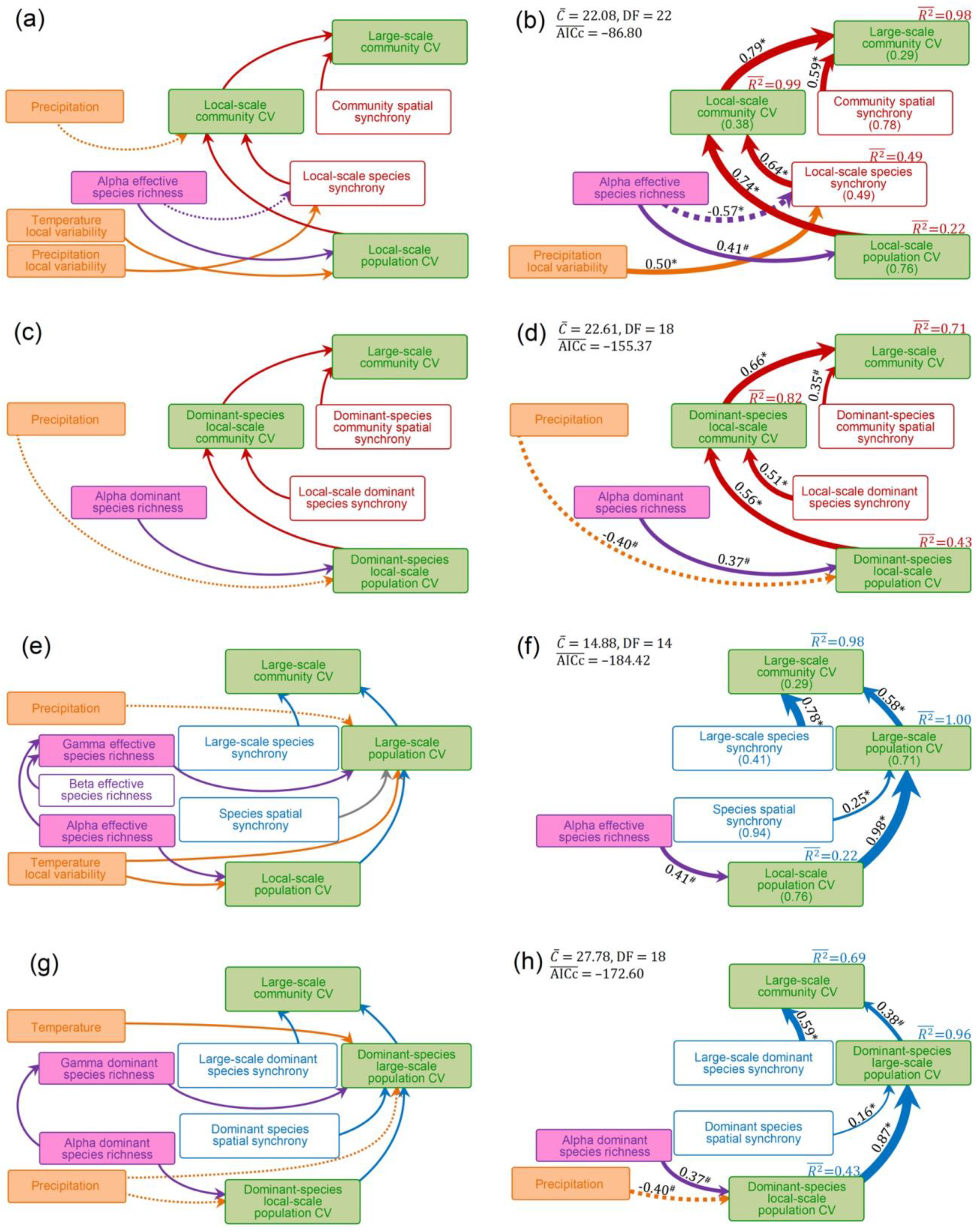
Initial (a, c, e and g) and final (b, d, f and h) structural equation models (SEMs) relating the large-scale community coefficient of variation (CV, inverse of temporal stability) to CVs and synchronies (inverse of asynchronies) at lower hierarchical levels of ecological organization and to species diversity indices estimated with all species (a, b, e and f) and only dominant species (c, d, g and h), as well as climatic factors. These SEMs separately considered the upscaling pathways of aggregating local-scale community (pathway I, a, b, c and d) or organizing large-scale population (pathway II, e, f, g and h). In initial SEMs (a, c, e and g, which can also be found in Figure 4–figure supplement 3b–3c and Figure 4–figure supplement 4b–4c), colored and grey arrows respectively represent significant (or marginal significant) and non-significant paths and solid and dashed arrows respectively represent positive and negative paths (see Figure 4–figure supplement 1 for detail). In final SEMs (b, d, f and h), solid and dashed colored arrows respectively represent examined positive and negative paths (Figure 4– source data 1–2), which have also been scaled in relation to the strength of the relationship with numbers showing the mean values the standardized path coefficients. 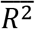 values are mean values of proportion of variance explained by dependent variables in the model. In addition, in the final SEM for all-species measures (b and f), mean values of CVs and synchronies have been shown in brackets. The significance level of each path is indicated by * when *P* < 0.05 or # when *P* < 0.10 (see Materials and Methods for details). Diagrams of final SEMs combining different upscaling pathways can be found in Figure 4. Dataset, code and relevant results can also be found in Figshare https://doi.org/10.6084/m9.figshare.16903309.

**Figure 6–figure supplement 1.**
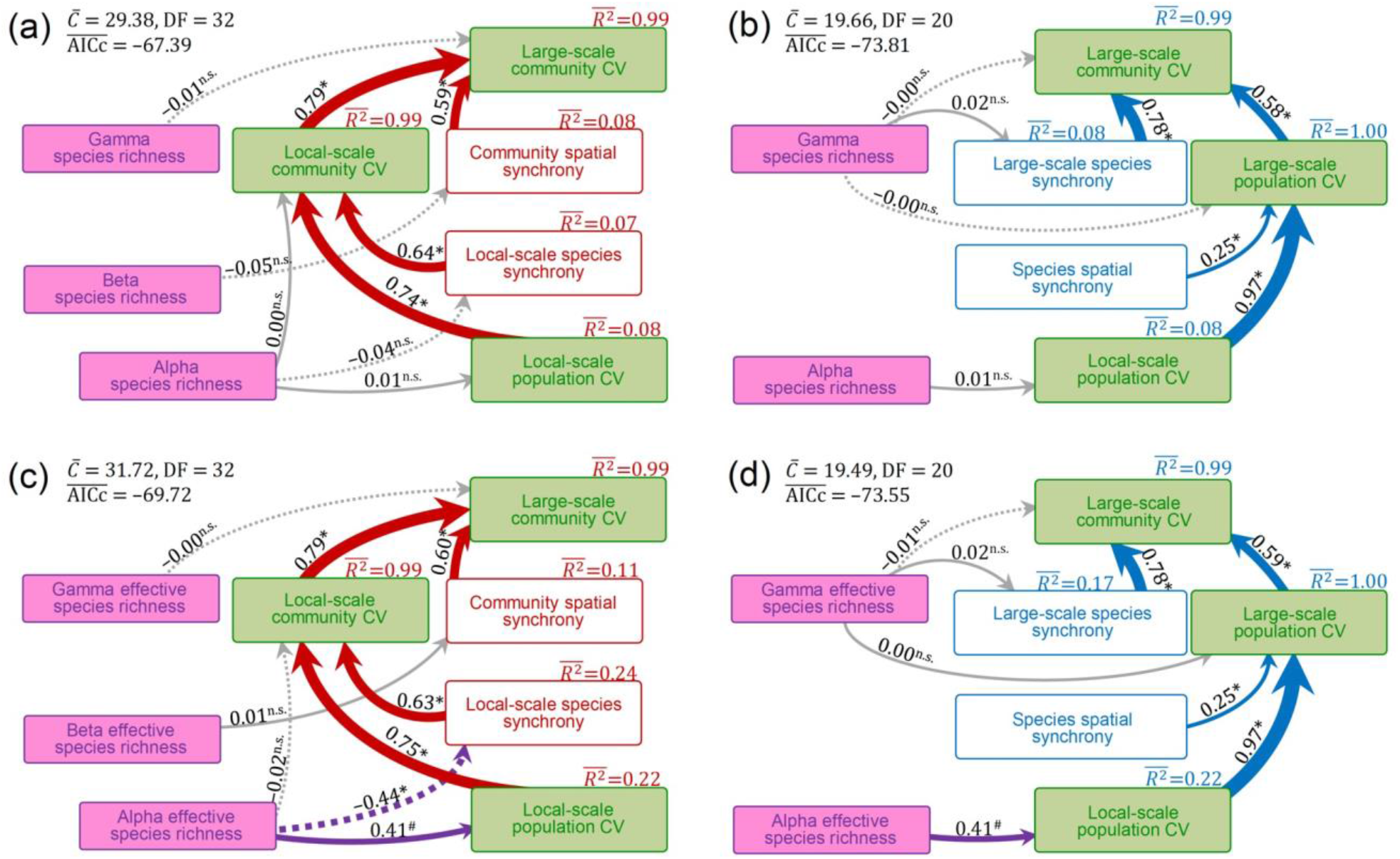
Diagrams of structural equation models (SEMs) examining the theoretically proposed impacts of species diversity (species richness, a–b, and effective species richness, c–d) on the large-scale community coefficient of variation (CV, inverse of temporal stability) and its hierarchical components with separately considering the upscaling pathways of aggregating local-scale communities (pathway I, a and c) and organizing large-scale populations (pathway II, b and d). Details can also be found in Figure 6–source data 1. Colored and grey arrows represent significant (or marginal significant) and non-significant paths, respectively. Solid and dashed arrows, respectively, represent examined positive and negative paths (Figure 6– source data 1). Arrows have also been scaled in relation to the strength of the relationship with numbers showing the mean values the standardized path coefficients. The significance level of each path is indicated by * when *P* < 0.05, # when *P* < 0.10 or n.s. (non-significant) when *P* > 0.10 (see Materials and Methods for details). Diagrams of SEMs combining different upscaling pathways can be found in Figure 6. Dataset, code and relevant results can also be found in Figshare https://doi.org/10.6084/m9.figshare.16903309.

## Supplementary file 1

### Time series of plant species biomass in each surveyed site

**Supplementary file 1–Figure 1.**
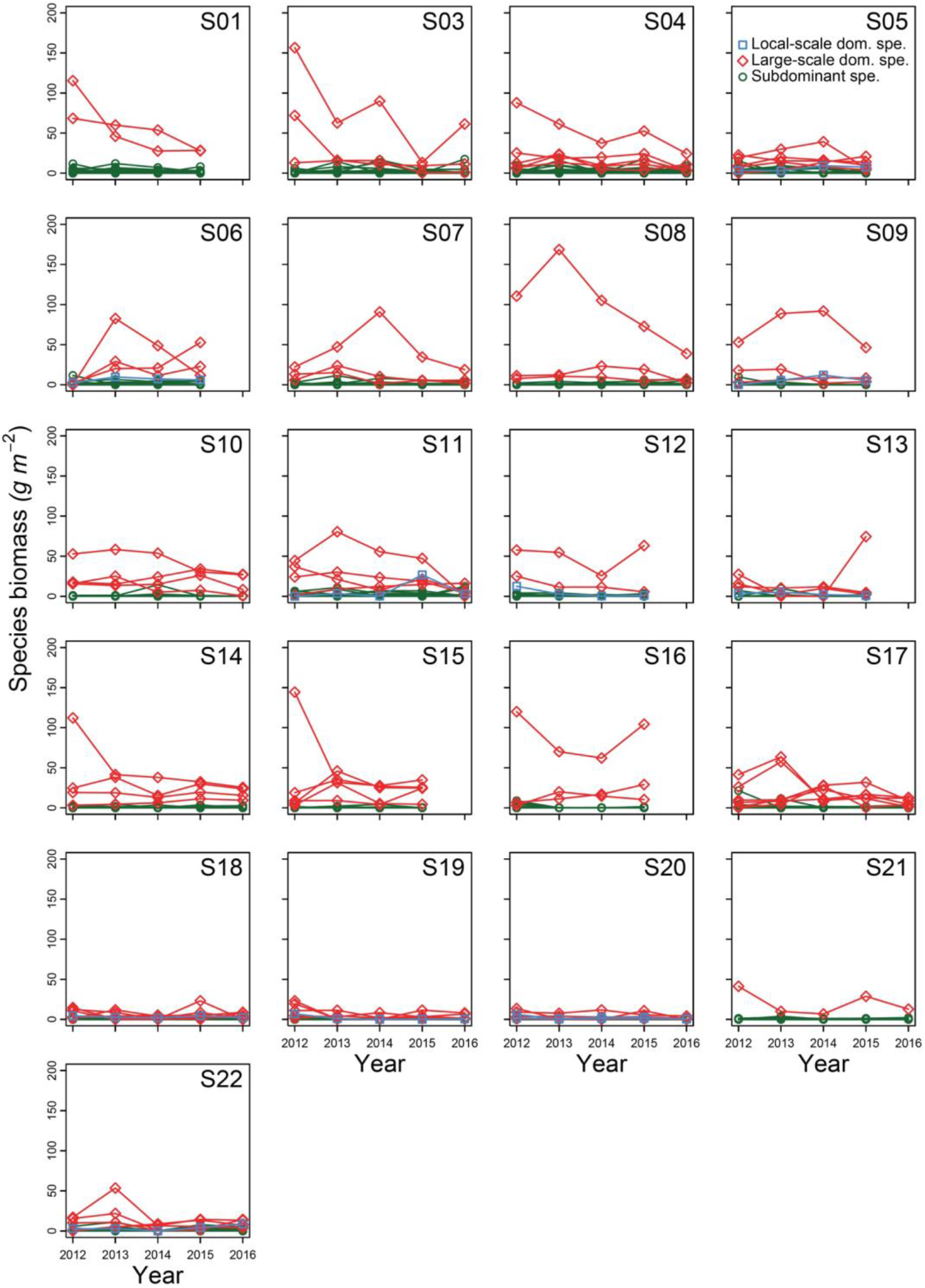
Time series of plant species biomass in each surveyed site. Blue squares and lines represent species that only characterized as dominant species in local-scale communities. Red diamonds and lines represent species characterized as dominant species in local-scale communities and can also be characterized as dominant species when aggregating into large-scale communities. Green circles and lines represent subdominant species. It showed that most dominant species of local-scale communities can be defined as dominant species of large-scale communities, with only few exceptions. In addition, these species have higher productivity than others roughly all the time and are constantly exist in surveyed sites. Dataset, code and relevant results can also be found in Figshare https://doi.org/10.6084/m9.figshare.16903309.

### Supplementary file 2

#### Mathematical derivation for partitioning temporal stability and synchrony across ecological hierarchies into dominant and subdominant species groups

Here, we introduce mathematical derivations used to partition large-scale community temporal stability and its hierarchical components into dominant and subdominant species groups. These derivations based on previous theoretical investigations of (temporal) coefficient of variation (CV, inverse of temporal stability) and synchrony (inverse of asynchrony) across ecological hierarchies (Thibaut and Connolly, 2013; Wang et al., 2019; Wang and Loreau, 2016, 2014). Briefly, these investigations have shown that local-scale population CV can be upscaled to that of large-scale community with two alternative pathways I or II. In the first upscaling pathway (pathway I), local-scale populations organize into local-scale communities, and then, local-scale communities aggregating into a large-scale community (Wang et al., 2019; Wang and Loreau, 2016, 2014) (Figure 1b). In another upscaling pathway (pathway II), local-scale populations scale up to large-scale populations, and then, large-scale populations organizing into a large-scale community (Wang et al., 2019; Wang and Loreau, 2016, 2014) (Figure 1b). In each upscaling pathway, synchrony at lower organization level or spatial scale determines the proportion of CV upscaled to higher organization level or spatial scale (Wang et al., 2019; Wang and Loreau, 2016, 2014). In the upscaling pathway of aggregating local-scale communities (pathway I), local-scale population CV firstly upscales to local-scale community CV with local-scale species synchrony measuring the proportion of CV transformed to local-scale community (Loreau and de Mazancourt, 2008; Thibaut and Connolly, 2013; Wang et al., 2019; Wang and Loreau, 2016, 2014). Subsequently, local-scale community CV upscales to large-scale community CV with community spatial synchrony measuring the proportion (Wang et al., 2019; Wang and Loreau, 2016, 2014). In the upscaling pathway of organizing large-scale populations (pathway II), local-scale population CV first upscales to large-scale population CV with species spatial synchrony measuring how much CV has been upscaled, then, upscaling to the large-scale community CV with large-scale species synchrony measuring the proportion (Wang et al., 2019). Descriptions of these terms can be found in Table 1.

In the following part, we only introduce methods partitioning CVs and synchronies across ecological hierarchies into dominant (relative species abundance > 5%, see Supplementary file 1 for details) and subdominant species groups without repeating previous theoretical derivations relating them across different hierarchies but recommended readers to these works for further details (Loreau and de Mazancourt, 2008; Thibaut and Connolly, 2013; Wang et al., 2019; Wang and Loreau, 2016, 2014). We used superscripts *P* and *C* to designate the quantities of population level and community level, superscripts *L* and *A* the quantities of localities (e.g. local-scale communities) and an aggregation of multiple localities (e.g. large-scale communities aggregating multiple local-scale communities), and superscript *P→C* and *L→A* the organization of populations into communities and aggregation of local-scale units into large scales. Symbols and descriptions used in the following partitions can be found in Table 1.

We consider a large-scale community reached a stationary state, which includes *M* localities (e.g. sites or local-scale communities) and *N* species. This large-scale community can be described with a matrix of (temporal) mean species abundance with elements *u^P,L^*(*i*, *k*), i.e. the mean abundance of species *k* in locality *i*, and a (temporal) variance–covariance matrix of species abundances with elements *v^P,L^*(*ij*, *kl*) = *cov*(*u^P,L^*(*i*, *k*), *u^P,L^*(*j*, *l*)), i.e. the covariance between abundances of species *k* in locality *i* and species *l* in locality *j*. In addition, we introduce two matrixes, *d^P^* and *s^P^*, to represent the dominant and subdominant species of the large-scale community, respectively. For the *d^P^*, it has *M* rows and *N* columns, representing numbers of localities and species of the large-scale community, and has elements *d^P^*(*i*, *k*), i.e. the *k*th species of the *i*th locality, which is set to 1 if the *k*th species is a dominant species at the large scale, otherwise, 0. Similar procedure is used to conduct the *s^P^*, in which, subdominant species are set to 1, otherwise, 0.

#### Supplementary file 2A. Partitioning local-scale population CV into dominant and subdominant species groups

The local-scale population CV (*CV^P,L^*) is defined as the weighted average local-scale population CV, which can be described as follows (Thibaut and Connolly, 2013; Wang et al., 2019; Wang and Loreau, 2016, 2014):

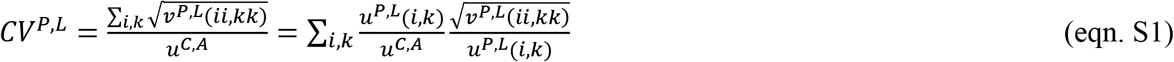

We rewrite this equation with introduced two matrixes (*d^P^*(*i*, *k*) and *s^P^*(*i*, *k*)) to separate the local-scale population CV (*CV^P,L^*) into its dominant 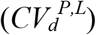 and subdominant 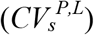 species group components, which has the following description:

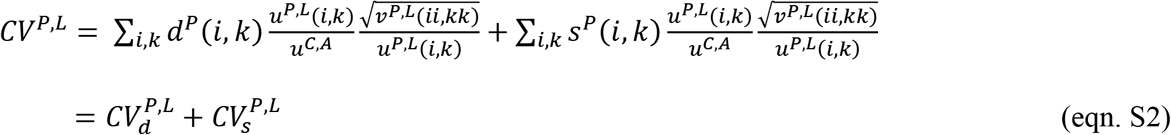

#### Supplementary file 2B. Partitioning local-scale species synchrony into dominant and subdominant species groups

The local-scale species synchrony (*φ^P→C,L^*) is defined as the weighted average synchronous dynamics among populations of different species within local-scale communities, which has the following description (Wang et al., 2019; Wang and Loreau, 2016, 2014):

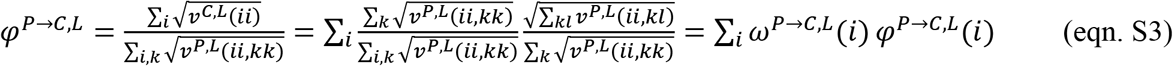

where *ω^P→C,L^*(*i*) and *φ^P→C,L^*(*i*) are the contribution of local-scale population variance of the *i*th community to the sum of variance of all species local-scale populations within the large-scale community and synchronous dynamics among local-scale populations of different species within the *i*th local-scale community (i.e. species synchrony of the *i*th local-scale community, Loreau & de Mazancourt 2008), respectively. We can rewrite *φ^P→C,L^*(*i*) with *d^P^*(*i*, *k*) and *s^P^*(*i*, *k*), which has the following description:

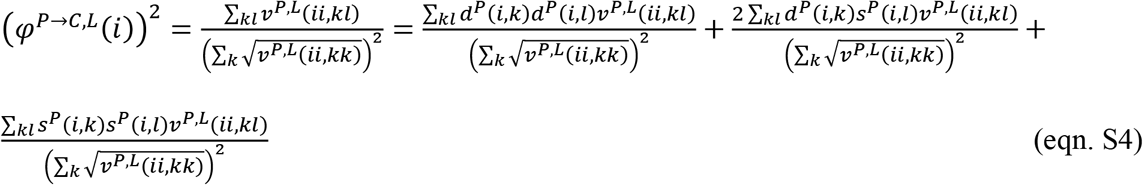

We defined the first term of the right-hand side of the eqn. S4 as the dominant-species local-scale species synchrony of the *i*th local-scale community (Wang et al., 2020), which has the following description:

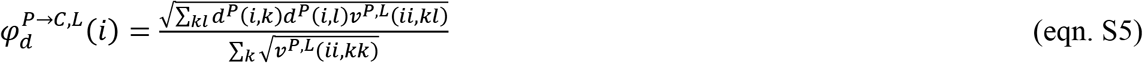

Then, using above description, we defined the dominant-species local-scale species synchrony of the large-scale community 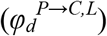, i.e. an aggregation of multiple local-scale communities, as the follows:

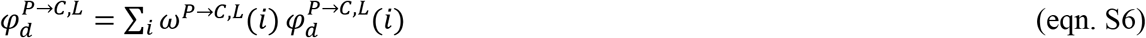

Referenced to the definition of local-scale community CV, *CV^C,L^ = φ^P→C,L^ × CV^P,L^* (Wang et al., 2019; Wang and Loreau, 2016, 2014), we defined the dominant-species local-scale community 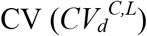 as follows:

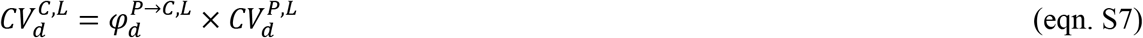

#### Supplementary file 2C. Partitioning community spatial synchrony into dominant and subdominant species groups

The community spatial synchrony (*φ^C,L→A^*) defined as the weighted average synchronous dynamics among spatially separated local-scale communities, which has the following description (Wang et al., 2019; Wang and Loreau, 2016, 2014):

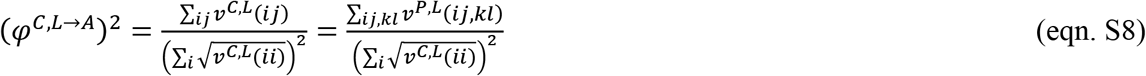

Using *d^P^*(*i*, *k*) and *s^P^*(*i*, *k*) mentioned above, we partitioned community spatial synchrony into dominant 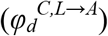, subdominant species groups 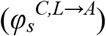 and synchronous dynamic between them 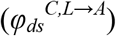 with the following description:

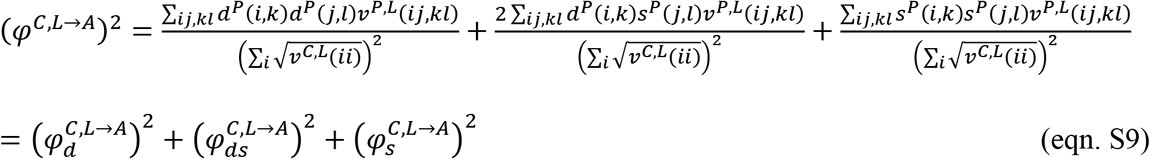

Referenced to the definition of large-scale community CV with the upscaling pathway of aggregating local-scale communities (pathway I), *CV^C,A^* = *φ^C,L→A^ × CV^C,L^* (Wang et al., 2019; Wang and Loreau, 2016, 2014), we defined the dominant-species large-scale community CV with this upscaling pathway 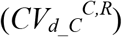 as follows:

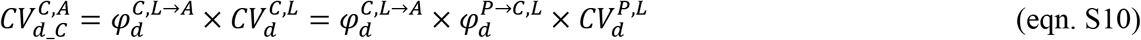

#### Supplementary file 2D. Partitioning species spatial synchrony into dominant and subdominant species groups

The species spatial synchrony (*φ^P,L→A^*) is defined as the weighted average synchronous dynamics among spatially separated local-scale populations of same species, which has the following description (Wang et al., 2019):

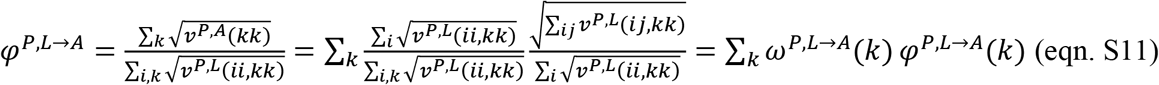

where *ω^P,L→A^*(*k*) and *φ^P,L→A^*(*k*) are the contribution of population variance of the *k*th species to that of all species within the large-scale community and synchrony within the *k*th species among sites, respectively. We can rewrite *φ^P,L→A^*(*k*) with *d^P^*(*i*, *k*) and *s^P^*(*i*, *k*), which has the following description:

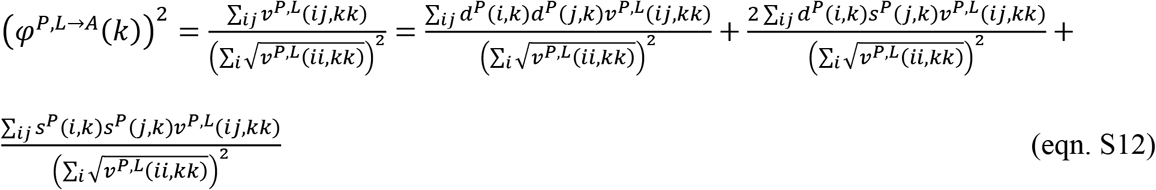

We defined the first term of the right-hand side of above equation as the species spatial synchrony of the *k*th (dominant) species, which has the following description:

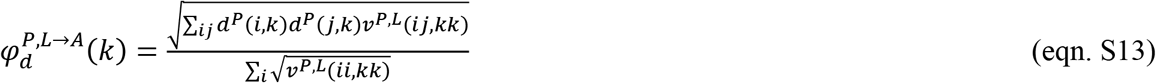

Then, using above description, we defined the dominant species spatial synchrony 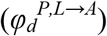 as the follows:

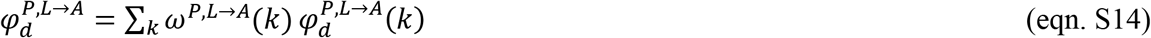

Referenced to the definition of large-scale population CV, *CV^P,A^* = *φ^P,L→A^ × CV^P,L^* (Wang et al., 2019), we defined the dominant-species large-scale population 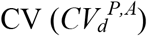 as follows:

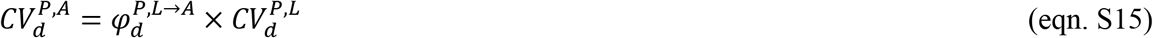

#### Supplementary file 2E. Partitioning large-scale species synchrony into dominant and subdominant species groups

The large-scale species synchrony (*φ^P→C,A^*) is defined as the weighted average synchronous dynamics among large-scale populations of different species, which has the following description (Wang et al., 2019):

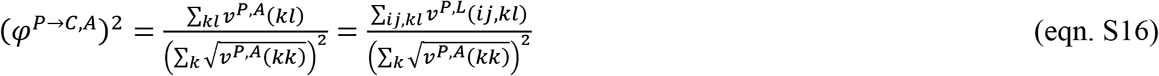

Here, *v^P,A^*(*kl*) is the covariance between *k* and *l* large-scale populations. We partitioned the large-scale species synchrony into dominant 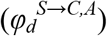, subdominant species groups 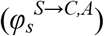 and synchronous dynamic between them 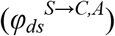 using introduced *d^P^*(*i*, *k*) and *s^P^*(*i*, *k*) with the following description:

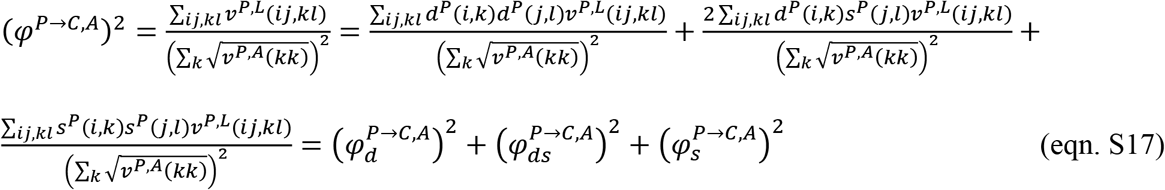

Referenced to the definition of large-scale community CV with the upscaling pathway of organizing large-scale populations (pathway II), *CV^C,A^* = *φ^P→C,A^ × CV^P,A^* (S. Wang et al., 2019), we defined the dominant-species large-scale community CV with this upscaling pathway 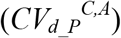 as follows:

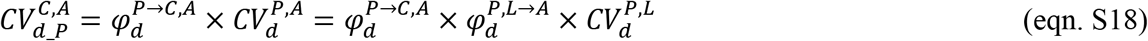

#### Supplementary file 2F. Comparing dominant-species large-scale community CVs estimated with two alternative upscaling pathways

Based on recent theoretical study (S. Wang et al., 2019), the large-scale community CV can be upscaled by aggregating local-scale communities 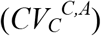 or organizing large-scale populations 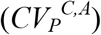, which have the following descriptions:

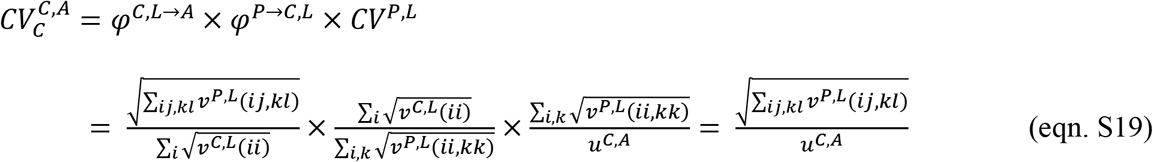

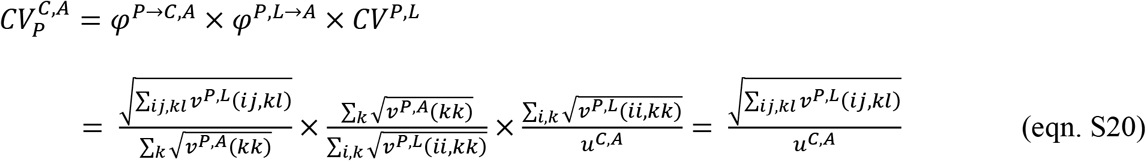

These descriptions (eqn. S19 and S20) showed that the large-scale community CV estimated with two different upscaling pathways are equal to each other.

In the following part, we explain why the dominant-species large-scale community CV estimated with two different upscaling pathways are not equal to each other (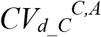 for estimated via aggregating local-scale communities, pathway I, and 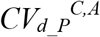 for estimated via organizing large-scale populations, pathway II). The dominant-species large-scale community CV estimated by aggregating local-scale communities 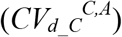 has the following description:

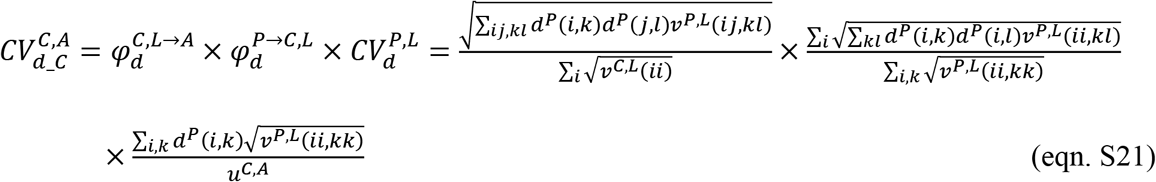

The dominant-species large-scale community CV estimated by organizing large-scale populations 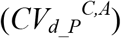 has the following description:

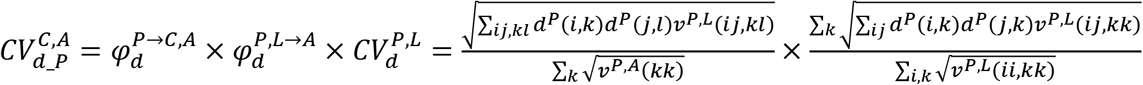

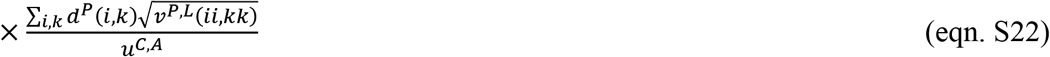

Owing to these two equations have either same terms or different terms 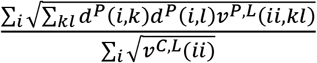 in eqn. S21 and 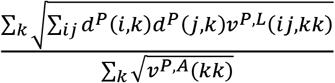 in eqn. S22), the dominant-species large-scale community CV estimated with two different upscaling pathways should be well correlated but not totally same. For the denominators of these two different terms, they are sum of local-scale community variances and sum of large-scale population variances. For the numerators of them, they are sum of variance (and covariance) of different dominant species within same local communities and sum of variances (and covariance) of same dominant species across different local communities. These differences reflect that dominant-species large-scale community CVs estimated via aggregating local-scale communities 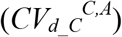 and via organizing large-scale populations 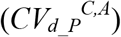 focus on different dominant species within same local-scale communities and same dominant species across different local-scale communities, respectively. Owing to the potential difference, we separately reported them (Supplementary file 5–Figure1a–b). It is also need to note that the different terms in eqn. S21 and eqn. S22 can be same when considering all species. This is because, in this case, they become to 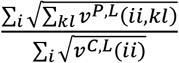 and 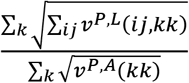, and both of them are equal to 1, resulting in same large-scale community CV estimated with all species using different upscaling pathways.

## Supplementary file 3

### Impacts of spatial distance on coefficients of variation and synchronies across spatial scales

**Supplementary file 3–Figure 1.**
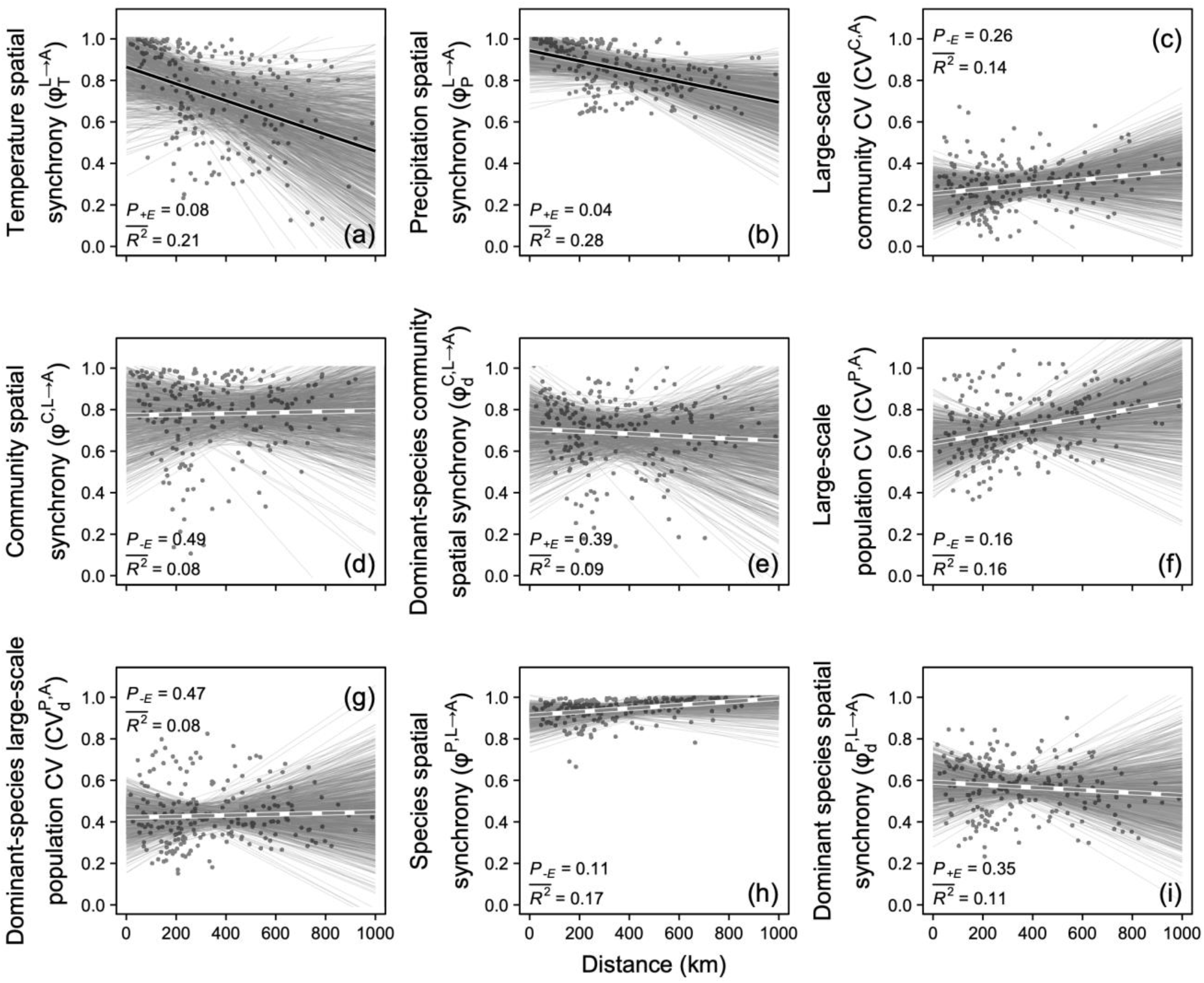
Spatial synchronies of temperature (a) and precipitation (b), large-scale community coefficient of variation (CV, inverse of temporal stabilities, c) and all-species and dominant-species estimates of community spatial synchrony (inverse of asynchrony, d and e), large-scale population CV (f and g) and species spatial synchrony (h and i) in relation to distance. Solid black lines represent significant (*P* < 0.05) and marginally significant (*P* < 0.10) and dashed grey line represents non-significant (*P* > 0.10) relationships (see Materials and Methods for details). Symbols and descriptions can be found in Table 1. Dataset, code and relevant results can also be found in Figshare https://doi.org/10.6084/m9.figshare.16903309.

## Supplementary file 4

### Results of general linear models examining impacts of species diversity on large-scale community coefficient of variation and its hierarchical components

**Supplementary file 4–Table 1.** General linear models for relating coefficients of variation (CVs, inverse of temporal stability) across organization levels and spatial scales to their hierarchical components and species diversity indices. Reported are the mean value of the estimated slope parameter 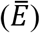 and its 10% and 90% quantiles (*QE_10_* and *QE_90_*), the mean values of the explanatory power 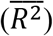, the proportion of *E* < 0 (*P_−E_*) when 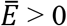 (or the proportion of *E* > 0, *P_+E_*, when 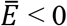), and the mean proportion of variance explained by the variables 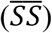. All these statistics are based on 1000 random splits of the dataset into ten large-scale communities each time. Dataset, code and relevant results can also be found in Figshare https://doi.org/10.6084/m9.figshare.16903309.

| Independent variable | $\bar{E}$ | $QE_{10}$ | $QE_{90}$ | $P_{-E}$ or $P_{+E}$ | $\bar{SS}$ |
| --- | --- | --- | --- | --- | --- |
| **Model 1: Large-scale community CV ~ Community spatial synchrony + Local-scale community CV + Gamma species richness**<br>$\bar{R}^2 = 0.99$ | | | | | |
| Community spatial synchrony | 0.586 | 0.356 | 0.838 | 0.000 | 0.417 |
| Local-scale community CV | 0.789 | 0.580 | 1.027 | 0.000 | 0.565 |
| Gamma species richness | -0.007 | -0.076 | 0.055 | 0.447 | 0.002 |
| **Model 2: Large-scale community CV ~ Community spatial synchrony + Local-scale community CV + Gamma effective species richness**<br>$\bar{R}^2 = 0.99$ | | | | | |
| Community spatial synchrony | 0.599 | 0.365 | 0.853 | 0.000 | 0.417 |
| Local-scale community CV | 0.790 | 0.588 | 1.023 | 0.000 | 0.565 |
| Gamma effective species richness | -0.004 | -0.071 | 0.066 | 0.443 | 0.003 |
| **Model 3: Large-scale community CV ~ Community spatial synchrony + Local-scale community CV + Beta species richness**<br>$\bar{R}^2 = 0.98$ | | | | | |
| Community spatial synchrony | 0.588 | 0.358 | 0.848 | 0.000 | 0.417 |
| Local-scale community CV | 0.788 | 0.584 | 1.028 | 0.000 | 0.565 |
| Beta species richness | 0.004 | -0.057 | 0.067 | 0.464 | 0.002 |
| **Model 4: Large-scale community CV ~ Community spatial synchrony + Local-scale community CV + Beta effective species richness**<br>$\bar{R}^2 = 0.98$ | | | | | |
| Community spatial synchrony | 0.593 | 0.359 | 0.850 | 0.000 | 0.417 |
| Local-scale community CV | 0.792 | 0.584 | 1.028 | 0.000 | 0.565 |
| Beta effective species richness | 0.008 | -0.060 | 0.073 | 0.417 | 0.002 |
| **Model 5: Large-scale community CV ~ Local-scale species synchrony + Local-scale population CV + Gamma species richness**<br>$\bar{R}^2 = 0.72$ | | | | | |
| Local-scale species synchrony | 0.500 | 0.211 | 0.756 | 0.014 | 0.337 |
| Local-scale population CV | 0.568 | 0.261 | 0.827 | 0.022 | 0.345 |
| Gamma species richness | 0.017 | -0.230 | 0.269 | 0.458 | 0.038 |
| **Model 6: Large-scale community CV ~ Local-scale species synchrony + Local-scale population CV + Gamma effective species richness**<br>$\bar{R}^2 = 0.72$ | | | | | |
| Local-scale species synchrony | 0.510 | 0.180 | 0.826 | 0.035 | 0.337 |
| Local-scale population CV | 0.564 | 0.216 | 0.867 | 0.040 | 0.345 |
| Gamma effective species richness | 0.021 | -0.341 | 0.401 | 0.487 | 0.041 |
| **Model 7: Large-scale community CV ~ Local-scale species synchrony + Local-scale population CV + Beta species richness**<br>$\bar{R}^2 = 0.72$ | | | | | |
| Local-scale species synchrony | 0.503 | 0.209 | 0.786 | 0.023 | 0.337 |
| Local-scale population CV | 0.574 | 0.271 | 0.836 | 0.017 | 0.345 |
| Beta species richness | -0.021 | -0.274 | 0.277 | 0.441 | 0.036 |
| **Model 8: Large-scale community CV ~ Local-scale species synchrony + Local-scale population CV + Beta effective species richness**<br>$\bar{R}^2 = 0.72$ | | | | | |
| Local-scale species synchrony | 0.510 | 0.217 | 0.785 | 0.017 | 0.337 |
| Local-scale population CV | 0.574 | 0.261 | 0.842 | 0.018 | 0.345 |
| Beta effective species richness | 0.034 | -0.261 | 0.352 | 0.449 | 0.041 |
| **Model 9:** *Local-scale community CV ~ Local-scale species synchrony + Local-scale population CV + Alpha species richness* |  |  |  |  |  |
| **$\overline{R^2} = 0.99$** |  |  |  |  |  |
| Local-scale species synchrony | 0.642 | 0.490 | 0.812 | 0.000 | 0.473 |
| Local-scale population CV | 0.736 | 0.588 | 0.900 | 0.000 | 0.518 |
| Alpha species richness | 0.003 | -0.041 | 0.046 | 0.454 | 0.001 |
| **Model 10:** *Local-scale community CV ~ Local-scale species synchrony + Local-scale population CV + Alpha effective species richness* |  |  |  |  |  |
| **$\overline{R^2} = 0.99$** |  |  |  |  |  |
| Local-scale species synchrony | 0.632 | 0.469 | 0.807 | 0.000 | 0.473 |
| Local-scale population CV | 0.746 | 0.593 | 0.909 | 0.000 | 0.518 |
| Alpha effective species richness | -0.016 | -0.085 | 0.052 | 0.381 | 0.001 |

## Supplementary file 5

### All-species measures of coefficients of variation and synchronies across spatial scales in relation to their dominant-species counterparts

**Supplementary file 5–Figure 1.**
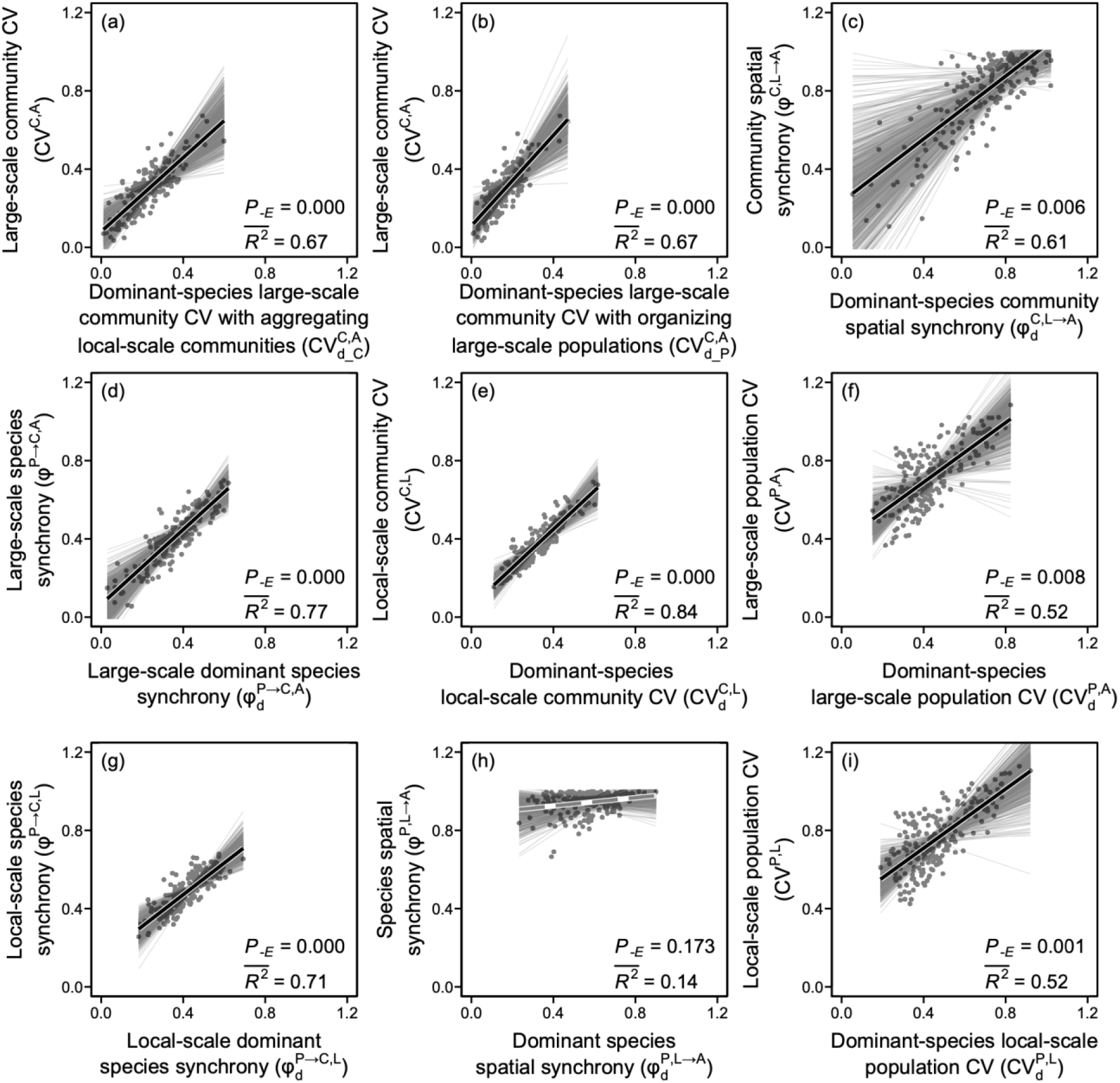
Coefficients of variation (CVs, inverse of temporal stabilities) and synchronies (inverse of asynchronies) across hierarchical levels of ecological organization in relation to their dominant-species counterparts. Solid black lines represent significant (*P* < 0.05) and marginally significant (*P* < 0.10) relationships and dashed grey line represents non-significant (*P* > 0.10) relationship (see Materials and Methods for details and Supplementary file 2F for estimating dominant-species large-scale community CV with upscaling pathways of aggregating local-scale communities, pathway I, and organizing large-scale populations, pathway II). Dataset, code and relevant results can also be found in Figshare https://doi.org/10.6084/m9.figshare.16903309.

